# LISA: A Case For Learned Index based Acceleration of Biological Sequence Analysis

**DOI:** 10.1101/2020.12.22.423964

**Authors:** Darryl Ho, Saurabh Kalikar, Sanchit Misra, Jialin Ding, Vasimuddin Md, Nesime Tatbul, Heng Li, Tim Kraska

## Abstract

Next Generation Sequencing (NGS) is transforming fields like genomics, transcriptomics, and epigenetics with rapidly increasing throughput at reduced cost. This also demands overcoming performance bottlenecks in the downstream analysis of the sequencing data. A key performance bottleneck is searching for exact matches of entire or substrings of short DNA/RNA sequence queries in a long reference sequence database. This task is typically performed by using an index of the reference - such as FM-index, suffix arrays, suffix trees, hash tables, or lookup tables.

In this paper, we propose accelerating this sequence search by substituting or enhancing the indexes with machine learning based indexes - called learned indexes - and present LISA (Learned Indexes for Sequence Analysis). We evaluate LISA through a number of case studies – that cover widely used software tools; short and long reads; human, animal, and plant genome datasets; DNA and RNA sequences; various traditional indexing techniques (FM-indexes, hash tables and suffix arrays) – and demonstrate significant performance benefits in a majority of them. For example, our experiments on real datasets show that LISA achieves speedups of up to 2.2 fold and 4.7 fold over the state-of-the-art FM-index based implementations for exact sequence search modules in popular tools bowtie2 and BWA-MEM2, respectively.

**Code availability:** LISA-based FM-index: https://github.com/IntelLabs/Trans-Omics-Acceleration-Library/tree/master/src/LISA-FMI

LISA-based hash-table: https://github.com/IntelLabs/Trans-Omics-Acceleration-Library/tree/master/src/LISA-hash

LISA applied to BWA-MEM2: https://github.com/bwa-mem2/bwa-mem2/tree/bwa-mem2-lisa.

## 1 INTRODUCTION

The latest high-throughput sequencers can read several terabases of DNA/RNA sequence data per day [1–3]. For example, a single Illumina Novaseq X Plus short read sequencer, launched in September 2022 [4], can sequence more than 128 genomes at 30x coverage in a 48-hour run generating 16 Terabases of data [1] at just $200 per genome [5]. The run sequences nearly 104 billion paired-ended reads, each of length 150 base-pairs. Among long read sequencers, a single PromethION 48 platform from Oxford Nanopore Technologies (ONT) can generate nearly 14 Terabases of data in a 72 hour run [3], generating reads of length up to a few million bases. The sequencing throughput is increasing and the cost is decreasing at a massive rate. Already today, a growing number of public and private sequencing centers with hundreds of NGS deployments are paving the way for population-scale genomics and transcriptomics applications. However, realizing this vision in practice heavily relies on building scalable systems for downstream sequence data analysis.

A fundamental step of downstream analysis is searching (a.k.a., sequence mapping) of millions of sequence queries (short or long reads) in a database of reference sequences (e.g., the human genome reference sequence consists of 3 billion bases). BWA-MEM (and its faster version BWA-MEM2) [6, 7], Bowtie2 [8], minimap2 [9] and STAR aligner [10] are among of the most widely used tools for sequence mapping. The key operation that has been shown to constitute a significant performance bottleneck during this mapping process is the search for *exact* matches of substrings of reads over the given reference sequence [6–8, 10–16]. This is typically done by building an index of the database to accelerate the search [6, 8, 10, 12–15]. Several data structures are proposed in the literature for the index — hash tables, FM-index based on Burrows Wheeler transform, suffix trees, suffix arrays, prefix tries, and prefix directed acyclic word graph. Out of these, hash tables (minimap2 [9], MAQ [17], SOAP [18]) and FM-index (Bowtie [12], Bowtie2 [8], SOAP2 [13], BWA [15], BWA-MEM2 [7]) are the most popular, while suffix trees were used by a few earlier tools (e.g., Mummer [19]).

Recently, in the databases domain, machine learning based index structures (a.k.a., learned indexes) have been shown to accelerate database queries [20]. In this work, we make a case for using learned indexes to accelerate sequence analysis; in particular, biological sequence search; and present LISA - Learned Indexes for Sequence Analysis (Figure 1).

**Figure 1:**
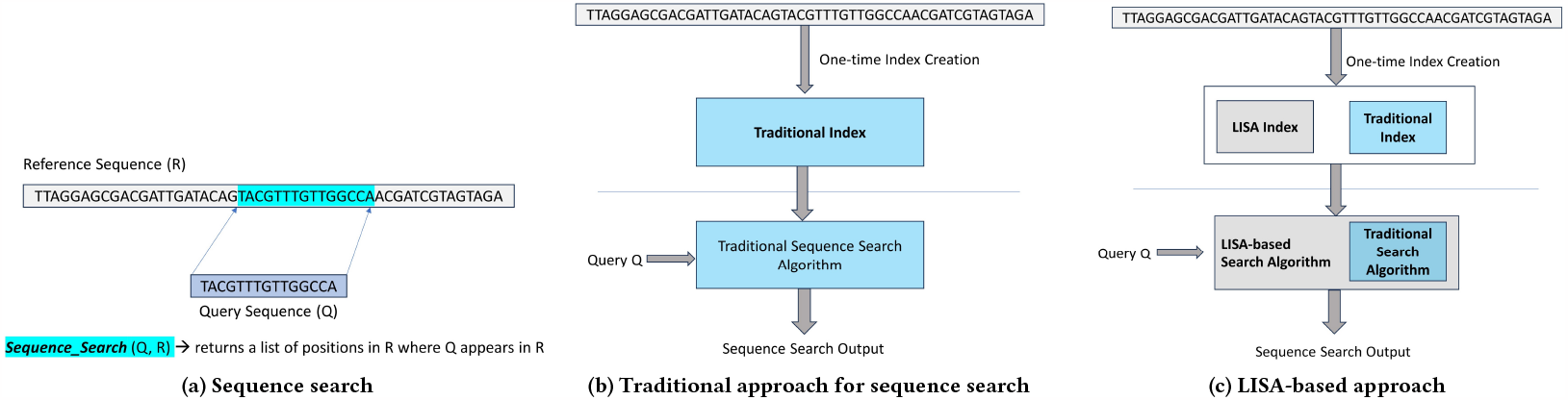
Our Proposed Approach. Searching of exact matches of entire or substrings of queries in a reference sequence is a fundamental task in computational biology. This task is traditionally done through an index of the reference sequence. We propose to enhance or substitute traditional indexes with machine learning based indexes.

Since we first introduced LISA in a preprint [21], multiple published papers [16, 22–24] have adopted this approach successfully to specific applications, thus, demonstrating its efficacy.

In this paper, we make a case for substituting or enhancing traditional indexes with learned indexes in a wider range of biological sequence search applications through a number of case studies. We present our results on using learned approach to enhance sequence search based on FM-index [25], which is an indexing technique employed by several tools, including BWA-MEM/BWA-MEM2 and Bowtie2. We also discuss how i) our prior work [16] accelerated the hash-tables based sequence search in minimap2, and ii) the sequence search in STAR aligner, that uses suffix arrays as indexing technique, can be performed more efficiently using an FM-index and thereby, can use our learned approach to enhance FM-Index.

In case of FM-index based exact sequence search, BWA-MEM2 and Bowtie2 use the following two key flavors and their variants: 1)*exact search*: search of matches of fixed length substrings of a read in the reference sequence and 2) *super-maximal exact match (SMEM) search*: for every position in the read, search of the longest exact match covering the position. Therefore, in the case studies presented in Section 2, we have focused on the variants of *exact search* and *SMEM search* that are used in these tools.

### 1.1 Sequence Search using LISA-enhanced FM-index

The FM-index implicitly represents the lexicographically sorted order of all suffixes of the indexed sequences. The key idea behind an FM-index is that, in the lexicographically sorted order of all suffixes of the reference sequence, all matches of a short DNA sequence (a.k.a., a “query”) will fall in a single region matching the prefixes of contiguously located suffixes. The performance of FM-index based algorithms is typically hampered by two issues. First, we need to process the query one letter at a time to find the the matching regions thus performing work that is *O* (*n*) in the size of the query. Second, the search algorithm suffers from poor data locality leading to frequent cache misses, thus degrading performance. Over the years, many improvements have been made to make the FM-index more efficient, leading to several state-of-the-art implementations that are highly cache-and processor-optimized [8, 12–15, 22, 26– 32]. Hence, it becomes increasingly more challenging to further improve FM-index based search without algorithmic improvements to reduce the work required.

LISA reduces the work required by enabling quick processing of several query letters at a time, thus, reducing the number of steps required to find the matching region of suffixes. The core idea behind applying LISA to FM-index based search is to speed up the process of finding the right region of suffixes in the FM-index by learning the distribution of suffixes in the reference. Recent work on learned index structures has introduced the idea that indexes are essentially models that map input keys to positions and, therefore, can be replaced by other types of models, such as machine learning models [20]. For example, a B-tree index maps a given key to the position of that key in a sorted array. Kraska et al. show that using knowledge of the distribution of keys, we can produce a learned model, that outperforms B-trees in query time and memory footprint [20]. The learned model predicts the position of the key in a sorted array and a range of positions in which the key is guaranteed to occur. If the key is not found in the predicted position, a last mile search is conducted in the provided range to find the key. Taking a similar perspective, the FM-index can be seen as a model that maps a given query sequence to the single region matching the prefixes of contiguously located suffixes. Therefore, it can be replaced with a learned model. One way to do this is to create a lexicographically sorted array of prefixes of fixed length, say *k*, of all suffixes of the reference sequence and using a learned model to search over that. However, this can only be used to search queries of length up to *k*. We propose a new index - IPBWT (Index Paired Burrows Wheeler Transform) - that is inspired by the last to first mapping of the FM-index to enable exact search of arbitrary length queries while processing a fixed number of letters at a time.

In the rest of the paper, we use LISA interchangeably to refer to both our proposed learned approach for sequence search and our library that contains implementations of some of the key LISA based algorithms for sequence search.

### 1.2 Our Contributions

We make the following contributions in this work.

- We introduce the idea of using learned indexes to accelerate biological sequence search.
- We demonstrate how *exact search* and *SMEM search* problems can be solved using learned indexes.
- We develop LISA-based implementations of various variants of *exact search* and *SMEM search* used in BWA-MEM2 and Bowtie2.
- We develop an architecture-optimized implementation of LISA through cache-optimizations, vectorization and multi-threading. We focus our efforts on the CPU, as that is the most widely available architecture for DNA sequence search.
- We demonstrate the benefits of LISA using multiple generations of server-grade CPUs for real Human, Asian Rice and Zebra Fish datasets. LISA achieves up to 2.2× and 13.7× speedups over the state-of-the-art implementations for *exact search* and *SMEM search*, respectively. Bowtie2 uses *exact search* and therefore, can be accelerated through LISA. Our LISA-based implementation achieves up to 4.7× speedup for sequence search (a.k.a. seeding) in BWA-MEM2.
- In the Discussion Section, we talk about how our prior work has used LISA to accelerate minimap2 and how LISA can be applied to other tools, e.g. STAR aligner, MUMmer, etc.

The rest of the paper is organized as follows. Section 2 presents key results, followed by discussion in Section 3 and conclusions in Section 4. Section 5 provide details of our methods including theoretical foundations. Supplementary section provides more details in terms of algorithms and results of evaluation of individual techniques and design choices.

## 2 RESULTS

We demonstrate the efficacy of LISA by comparing the throughput (million-reads/sec) with FM-index based *exact search, SMEM search*, and sequence search phase of BWA-MEM2.

### 2.1 System Configuration

We evaluate our solution on two generations of server-grade CPUs: single Intel^®^ Xeon^®^ Platinum 8280 processor and Intel^®^ Xeon^®^ Platinum 8480+ processor as detailed in Supplementary Table 1 and referred to as CLX and SPR, respectively, from here on. For multi-threaded runs, we use 2 threads per core to get the benefit of hyper-threads. Optimizing file I/O is beyond the scope of this paper. Therefore, we do not include file I/O time in any of our results.

### 2.2 Correctness

In all cases, we have verified that output of the LISA-based approach is identical to that of the traditional FM-index based approach.

### 2.3 Performance Evaluation for *Exact Search* and *SMEM Search*

For the baseline comparison, we use the Trans-Omics Acceleration Library (TAL) [32, 33], which provides the architecture optimized implementations for traditional FM-index based *exact search* and *SMEM search* [7, 32, *33]*.

*To establish TAL as an appropriate baseline, (i) for exact search*, we demonstrate in Supplementary Section A.4 that TAL is up to 4.2× faster than recently published Sapling [22], that is shown to be 2× faster than Bowtie; (ii) for *SMEM search*, we note that the optimized SMEM kernel from TAL is also used in BWA-MEM2 [7], a widely used architecture-optimized implementation of BWA-MEM [6].

#### 2.3.1 Datasets

We use three reference sequences - Human, Asian Rice, and Zebra Fish, as detailed in Supplementary Table 2. For each of these reference sequences, we use multiple real read datasets (H1-H5 for Human, A1-A3 for Asian Rice, and Z1-Z3 for Zebra Fish) downloaded from sequence read archive [34] (Supplementary Table 3). All of these read datasets consist of 5 million reads each. The read datasets for Asian Rice and Zebra Fish have reads of length 151. For Human reference, we use two types of reads datasets: H1-H3 contain of 151 length reads and H4-H5 are of 101 length. The older sequencing technologies produce reads of length 101, so we use these 101 length datasets to show the compatibility of LISA with the older sequencing technologies. For *exact search*, we use 22 length seeds generated from the read datasets as that is the default seed length used in Bowtie2. Seeds are the small fixed-sized substrings generated from a read sequence. For generating seeds, we followed the same strategy as Bowtie2 [8] and generated 50 million seeds for each of the read datasets.

#### 2.3.2 Exact Search

*Exact search* finds all end-to-end matches of the 22-length seeds. The traditional FM-index based search matches one base at a time against the reference sequence and therefore takes 22 steps for end-to-end matching of 22-length seed. LISA processes a whole 22-length seed in one shot and finds its matches in a single step. Figure 2 compares the throughput achieved by LISA and TAL for *exact search* on a single thread and single CPU across different reference sequences, read datasets. The x-axis represents the reference sequence and read datasets, and the throughput is shown on the y-axis. Observe that LISA outperforms TAL across all datasets, achieving 1.1 −2.2× higher throughput. As mentioned earlier, *exact search* is used in Bowtie2 and, therefore, our implementation can be used to accelerate Bowtie2.

**Figure 2:**
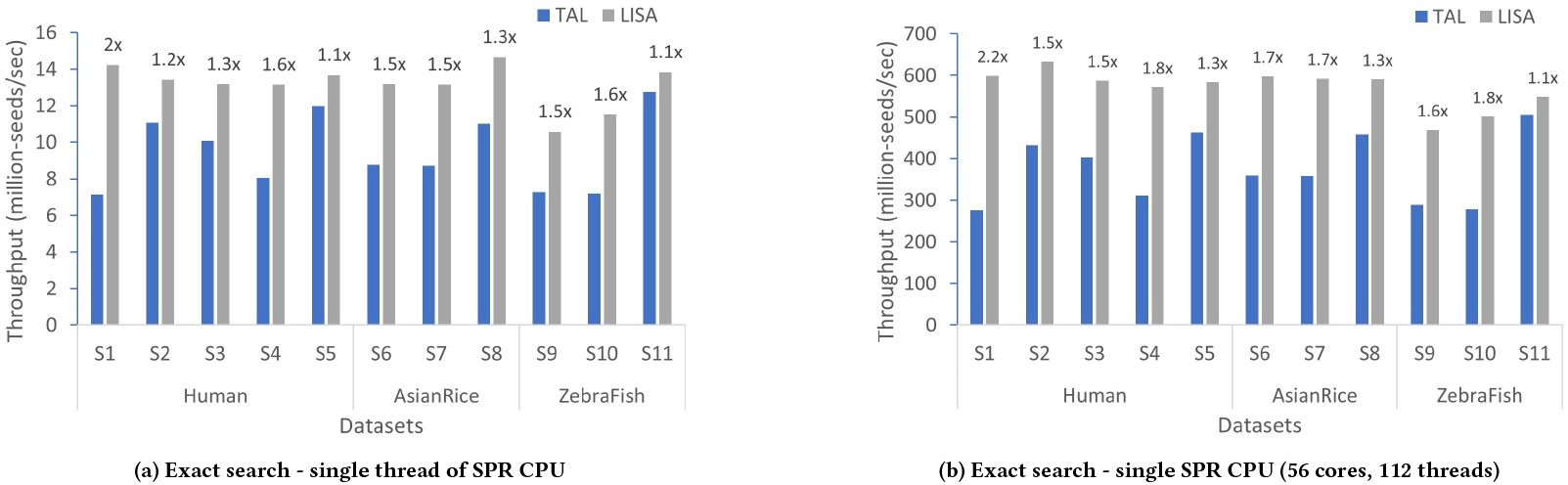
Performance comparison of LISA and TAL for *exact search* problem on a single thread and single SPR CPU with 56 cores. The speedup of LISA over TAL (the higher the better) for each dataset is shown on the top of the LISA bars.

#### 2.3.3 SMEM Search

Figure 3 presents the throughput comparison for *SMEM search* across different reference sequences, read datasets. LISA achieves 3.3 −13.7× higher throughput than TAL. On a single threaded execution on SPR, LISA achieves, on an average, 6.8× speedup over TAL across all datasets. For execution on an entire SPR CPU, LISA achieves, on an average, 4.8× speedup over TAL. This speedup is due to a combined effect of modification of the SMEM algorithm to enable usage of learned indexes and application of learned indexes.

**Figure 3:**
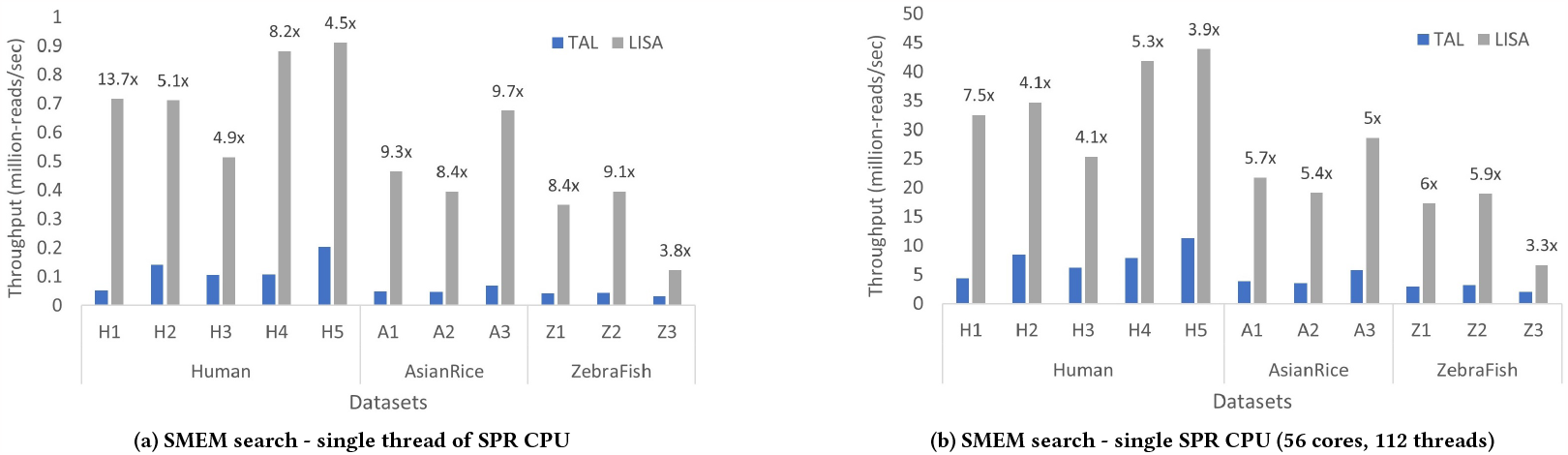
Performance comparison of LISA and TAL for *SMEM search* problem on a single thread and single of SPR CPU with 56 cores. The speedup of LISA over TAL for each dataset is shown on the top of the LISA bars.

Although LISA outperforms TAL across all datasets, the performance gain varies across datasets. The nature of reads and the reference sequences affect the overall performance gain. For instance, a read dataset with longer matching SMEMs is a better candidate for LISA than the one with the shorter matches. In Figure 3 the average length of the matches in H1 is 45, whereas in H3, the average length is 26. This correlates well with the performance gains.

### 2.4 Performance Evaluation for Seeding Phase of BWA-MEM2

Sequence search (a.k.a., seeding) in BWA-MEM2 consists of three modules that are variants of *exact search* and *SMEM search*. We developed LISA-based implementations of these modules and integrated them into BWA-MEM2. We compare the performance of LISA-based seeding against the seeding stage of BWA-MEM2 and also of BWA-MEME [23], a recently published approach to accelerate the seeding kernels of BWA-MEM2 using learned indexes.

#### 2.4.1 Datasets

We used the same 15 short read datasets that are used by the BWA-MEME paper [23] - out of which 6 datasets are from Illumina Platinum Genomes (length 101 bases) and 9 datasets are from 1,000 Genomes Project Phase 3 (length 150 bases). In addition to these 15 datasets, we also used four short read datasets with read length 250 to cover a wider range of read lengths. Supplementary Table 5 shows the details of all 19 datasets used in the experimentation.

#### 2.4.2 Performance Results

Figure 4 compares the speedup of BWA-MEM2, BWA-MEME, and LISA for paired-ended reads on two CPU generations. The x-axis shows the different datasets with their respective read lengths and the y-axis shows the corresponding throughputs. The speedup achieved by LISA over BWA-MEM2 across datasets is shown on top of LISA bars. Across all paired-ended reads LISA outperforms BWA-MEM2 and BWA-MEME and achieves up to 4.7× and 1.7× speedup compared to BWA-MEM2 and BWA-MEME, respectively. Section 6.2.2 explains how the different learned index based approaches used by LISA and BWA-MEME affect their respective performances. Supplementary Figure 1 shows that the throughput superiority of LISA holds for single-end reads as well. Even in this case, we have observed that the speedup difference across datasets is strongly correlated with length of matches and number of matches.

**Figure 4:**
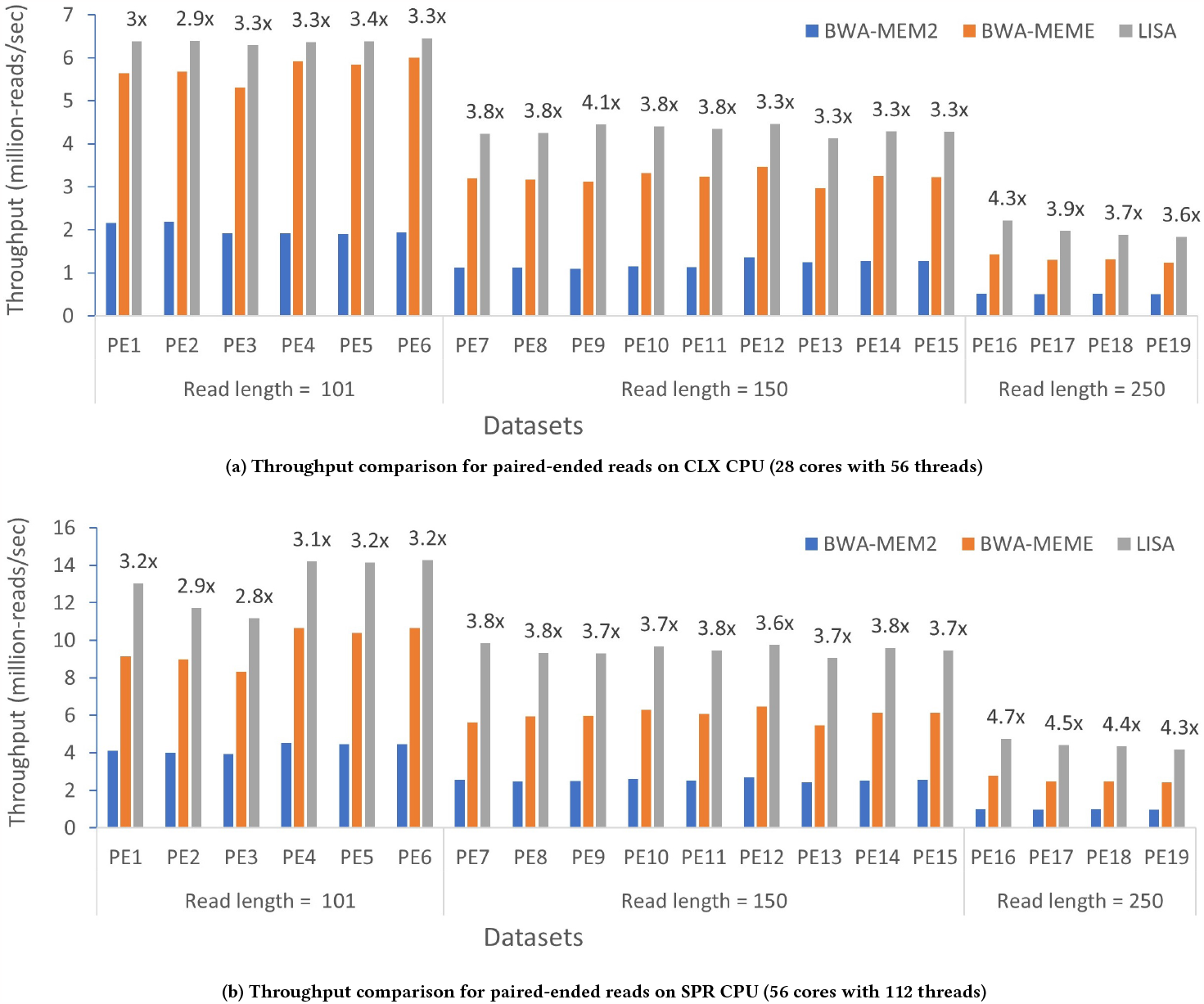
Performance comparison for the *seeding phase* of BWA-MEM2 on a single CLX CPU and SPR CPU for paired-ended datasets. The speedup of LISA over TAL is shown on the top of the LISA bars.

## 3 DISCUSSION

### 3.1 Wider Applicability of LISA

So far, we have demonstrate the efficacy of LISA on use-cases that involve short reads, sequences over DNA alphabet, and use FM-index as index. In this section, we discuss the applicability of LISA beyond such use-cases.

Minimap2 is a widely used tool for mapping long reads to a reference sequence. During the seeding phase, minimap2 uses a hash table-based minimizer search, where minimizers are short fixed-length substrings (k-mers) of the reference sequence. Minimap2 stores the minimizers in a hash table for faster lookup. For each input read, minimap2 extracts all the minimizers from the read and performs a hash lookup for the matching minimizers. In our prior work [16], we replaced hash table-based minimizer search with LISA-based search on a sorted list of minimizers which results in 3 −4× faster minimizer lookup. For details on implementation and design choices, refer to the paper [16].

In the case of sequences over RNA alphabet, STAR aligner is among the most widely used tool for mapping short RNA reads to a reference sequence. During its seeding phase, STAR uses a suffix array based index of the reference sequence to search for maximal mappable prefix (MMP) between the read and the reference sequence. MMP is similar to the maximal exact match (MEM) concept used in BWA-MEM2. FM-indexes have been shown to be more efficient that suffix arrays for finding MEMs. Therefore, MMPs can be searched more efficiently using an FM-index; and thereby; a LISA-based index of the reference sequence.

MUMmer is a popular tool for aligning genomes [19]. MUMmer uses maximal unique matches (MUM) as seeds to anchor the alignment between genomes. MUMs are the maximal (can not be extended on either side) matching sub-strings occurring only once in either of the genomes. MUMmer employs *generalized suffix tree* – suffix tree constructed from both the genome sequences. To find MUMs, a scan is made to the suffix tree to extract the internal nodes having 1 child from each genome. The sub-string from root to such internal nodes represents unique matches, the maximal among the unique matches represents MUMs. We propose the search for MUMs can be done using FM-index of the two genomes combined. Once constructed, the suffixes from the smaller genome can be searched in the index. A MUM is found in FM-index only if the SA-interval has two entries one from each genome. LISA, as shown for *SMEM search*, can be used to accelerate the suffix searches in the index.

The above examples demonstrate the following:

- LISA’s utility extends beyond exact and SMEM matches, encompassing various types of seed searches including MEMs, MMPs and minimizer searches.
- LISA can potentially serve as an efficient accelerator for sequence searches that have traditionally employed indexing methods, such as, Suffix Arrays/Trees, Hash Tables, and FM-index.
- LISA’s applicability is not restricted to short read DNA sequencing alone. Rather, it can be applied to long read sequencing as well as RNA sequencing.

### 3.2 Limitations

It is non-trivial to apply LISA to searching for inexact matches of a query sequence, where a few mismatches, insertions, and deletions are allowed. That is perhaps better done by processing the query one letter at a time. Moreover, in case of LISA-based *exact search* and *SMEM search* methods presented in this paper, LISA index requires larger memory compared to FM-index. For instance, for human genome of size 3 GB, FM-Index based solutions typically require 4.8 GB and 9.5 GB of index for *exact search* and *SMEM search*, respectively. LISA additionally requires 33 GB and 92 GB of index for the *exact search* and *SMEM search*, respectively. Therefore, it may not be a suitable technique for machines with smaller memory. However, with the latest CPUs that are typically accompanied by large memory, this is not a concern.

## 4 CONCLUSIONS AND FUTURE WORK

Using an index of a sequence database to search for sequence queries appears as a motif in many key areas in computational biology including genomics, transcriptomics, epigenetics and proteomics. In this paper, we presented LISA - a machine learning based approach to index a sequence database to accelerate sequence search. We demonstrated the benefits of our approach through several case studies of DNA/RNA sequence search covering popular software tools. In particular, we presented results for two flavors of DNA sequence search that appear in multiple software tools - *exact search* and *SMEM search*, and showed up to 2.2× and 13.7× speedup, respectively. As a proof of efficacy of LISA in real-world applications, 1) the *exact search* module can be used as is to accelerate Bowtie2, 2) LISA provides up to 4.7× speedup for the seeding stage in BWA-MEM2 and up to 4× speedup for minimizer lookup in minimap2. We also discussed how LISA can be used to accelerate STAR aligner and MUMmer. As future work, we plan to apply LISA to STAR aligner, explore application of LISA to other DNA, RNA and protein sequence search tools. We also plan to explore applications of learned indexes to many other problems in computational biology in which an index is created to accelerate search through a database.

## 5 METHODS

In this section, we define the notations and formally define the two DNA sequence search problems: *exact search* and *SMEM search*. For each problem, we briefly describe the prior work and established methods to solve these problems. We provide the background on learned indexes and explain how we use learned indexes to solve the two DNA sequence search problems. Subsequently, we show how we bring it all together to accelerate the seeding phase of BWA-MEM2. We end with the description of our hardware-aware efficient design and implementation.

For both exact search and SMEM search problems, an index of the reference sequence is required. Therefore, for all the algorithms in this section, the applicable index over the reference sequence is provided as input even if not explicitly stated, and we skip listing them for the sake of brevity.

### 5.1 Notation

We use upper case letters like *X, Y* to denote a DNA sequence, modeled as a string over the alphabet Σ = *{A,C,G,T}*, representing the four bases, and lower case letters like *x, y* to denote bases. Let |*X*| denote the length of *X, X* [*i*] denote the base at position *i*, and *X* [*i, j*] (*i*≤ *j* ) denote the substring *X* [*i*] *X* [*i* +1] *X* [*i*+2]… *X* [*j*] . We use *y*.*X* and *X* .*y* to represent prepending and appending, respectively, of base *y* to DNA sequence *X* and *X* .*Y* to denote the DNA sequence formed by concatenating sequences *X* and *Y* . We use *y*^*n*^ to denote a sequence composed of *n* contiguous copies of base *y*; e.g. *T* ^3^ is *TTT* .

We define 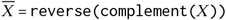, a reverse-complement of *X* where reverse(*X*) reverses the order of bases in *X* and complement(*X* ) maps each base of *X* to its complemented base using a mapping function {*A* → *T, C* → *G, G* → *C, T* → *A*}. For instance, given a DNA sequence 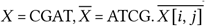 denotes the reverse-complement of substring *X* [*i*]*X* [*i* + 1]*X* [*i* + 2] · · · *X* [ *j* ] and 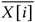 denotes the complemented base of *X* [*i*].

### 5.2 Exact Search Problem

Given a reference sequence *R* and a query sequence *Q*, the goal of exact sequence search is to find exact end-to-end matches of *Q* in *R*. More formally, an occurrence of *Q* is a position *p* in *R*, such that

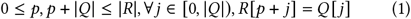

*Exact search* finds all such occurrences of *Q* in *R*. Typically, |*R*| ≈10^9^ bases; e.g., the length of the human genome is nearly 3 × 10^9^. On the other hand, |*Q*| typically ranges from a few tens to a few hundred bases; e.g., the default query length in Bowtie2 for *exact search* is 22.

### 5.3 FM-Index

The most widely used method to perform DNA sequence search is by creating an FM-index of the reference and searching for the query using the FM-index.

Fig. 5 depicts the construction of the FM-index for an example reference sequence *R*. First, we append *R* with the character $ ∉ Σ which is lexicographically smaller than all characters in Σ. Subsequently, we obtain all the rotations of *R* (Rotations(*R*)). The lexicographically sorted order of the rotations forms the BW-Matrix (Burrows Wheeler Matrix). The Burrows Wheeler Transform, a.k.a. BWT (*B*), is the last column of the BW-Matrix. The original positions in *R* of the first bases of these rotations constitute the suffix array (*S*).

**Figure 5:**
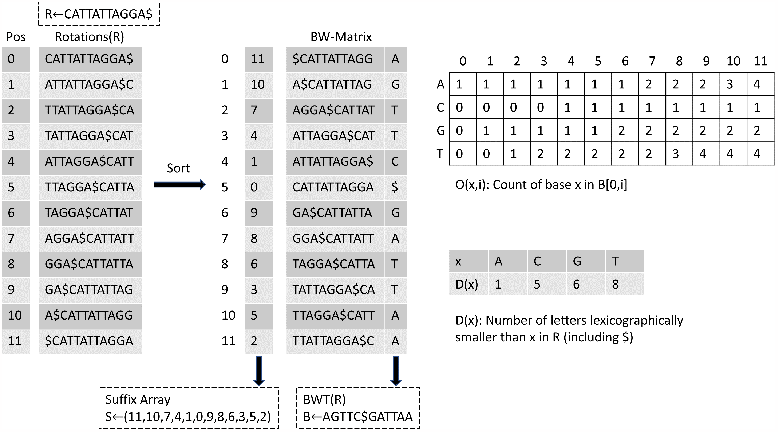
FM-index, *F* = ⟨*S, B, D, O*⟩ and BW-Matrix for sample reference sequence *R* ← ATACGAC$. The lexicographical ordering is $*< A < C < G < T* [32].

All the exact matches of a query can be found as prefixes of the rotations in the BW-Matrix. Since the BW-Matrix is lexicographically sorted, these matches are located in contiguous rows of the BW-Matrix. Therefore, for a query, all the matches can be represented as a range of rows of the BW-Matrix. This range is called the *SA (Suffix Array) interval* of the query. For example, in Fig. 5, the *SA interval* of query *“AC”* is [1, 3) . The values of the suffix array in the SA interval are 5 and 2. Indeed, the sequence *“AC”* is found at positions 5 and 2 in the reference sequence.

The FM-index is used to expedite search for the SA interval [25]. FM-index of a reference sequence *R, F* is be defined as the tuple ⟨*S, B, D, O*⟩, consisting of the suffix array *S* and the BWT *B*, as well as *D* and *O* data structures. *D* (*x*) is the count of bases in *R* [0, |*R*| −1] that are lexicographically smaller than *x*∈ Σ. *O* (*x, i*) is the count of occurrences of base *x* in *B* 0, *i* . Note that the BW-Matrix is not stored. For further information on the FM-index, see [25].

### 5.4 Exact Search using FM-Index

Figure 6 describes *exact search* of query in reference sequence. Exact search based on FM-index is performed using backward search algorithm (Alg. 1) [15]. The query is processed from the end to the beginning. For each value of *i*, the algorithm finds the SA interval [*l, r* ) of *Q* [*i*, |*Q* | − 1] in *R* by performing backward extension (line 5) on *SA interval* of *Q* [*i* + 1, |*Q* | − 1] by *Q* [*i*]. The backward extension is performed using the function *f* : (Σ, ℤ) → ℤ supported by the FM-index that takes a base *y* and an integer location *l* that is the lower bound of the *SA interval* of a DNA sequence *X*, and finds in *O* (1) time the lower-bound of the *SA interval* of a DNA sequence *y*.*X*, such that

**Figure 6:**
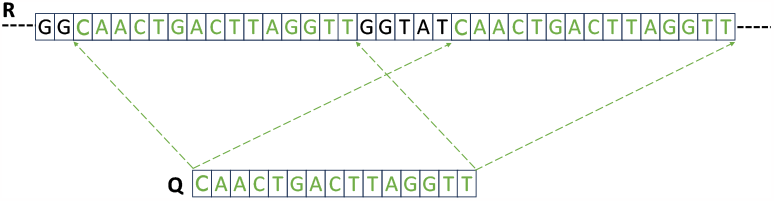
Exact query match between query and reference sequence. Complete query *Q* matches at two positions in reference sequence *R*.

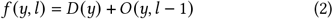

The width of the *SA interval* either shrinks or remains the same at each step. If the algorithm terminates before reaching the beginning of the query, *Q* does not occur in *R* (line 8). Otherwise, the range corresponds to the positions of exact occurrences of *Q* in *R* (line 9). Clearly, backward search algorithm requires at most | *Q* | backward extensions, each of which consumes constant time. Hence, the time complexity of the backward search algorithm is *O* (|*Q* |).

### 5.5 Learned Indexes

Recent work on learned index structures has introduced the idea that indexes are essentially models that map inputs to positions and, therefore, can be replaced by other types of models, such as machine learning models [20]. For example, a B-tree index maps a given key to the position of that key in a sorted array. Kraska et al. show that using knowledge of the distribution of keys can produce a learned model, e.g., the recursive model index (RMI), that outperforms B-trees in query time and memory footprint. Following [20], other learned index structures have been proposed [35–38].

Taking a similar perspective, the FM-index can be seen as a model that maps a given query sequence to the SA interval for that query sequence. Based on this insight, in LISA, we use knowledge of the distribution of subsequences within the reference sequence to create a learned index structure that enables faster exact and SMEM search.

### 5.6 Exact-LISA: Exact Search using LISA

In this section, we describe how LISA combines a new index data structure with learned indexes to improve the performance of exact search. We define our LISA index used for exact search as the tuple, *L* = ⟨ *F, I PBW T, RMI* ⟩, where *F* is the FM-index and we will define *I PBW T* and *RMI* below.

*Backward-search algorithm* performs exact search of a query *Q* using the FM-index by iterating through the query sequence in backwards order, one base at a time, thereby consuming *O Q* steps. The key idea of LISA is to iterate backwards through the query sequence in chunks of *K* bases at a time, so that exact search takes 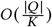 steps. Similar to *f*, we need to support a function, *f* ^*K*^ : (Σ^*K*^, ℤ) →ℤ, that takes in a length-*K* DNA sequence *Y* and the lower bound, *l*, of the *SA interval* of a DNA sequence *X* and returns the lower bound, *l* ^′^, of the *SA interval* of a DNA sequence *Y* .*X* . FM-Index can be redesigned to process *K* bases at a time by replacing Σ with Σ^*K*^ as the alphabet. However, it would require memory proportional to |Σ|^*K*^, and hence is infeasible.

#### 5.6.1 IPBWT

In LISA, to enable the search for K bases at a time, we designed a new data structure named **Index-Paired BWT (IPBWT)**. For each row of the BW-Matrix, there is an entry in IPBWT consisting of a ⟨Σ^*K*^, ℤ⟩ pair. The first part is the first *K* bases of the corresponding BW-Matrix row. The second part is the BW-Matrix location of the DNA sequence with the first *K* and the last *n* − *K* bases swapped. For two IPBWT entries ⟨*X, l*⟩ and ⟨*X* ^′^, *l* ^′^⟩, we define

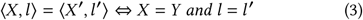

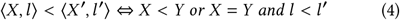

Note that, since the BW-Matrix is in a lexicographically sorted order, the IPBWT is also in a sorted order. Our desired function *f* ^*K*^ : (Σ^*K*^, ℤ) →ℤ is now equivalent to finding the lower bound location of an input ⟨*Y, l*⟩ in the IPBWT; i.e.; the first entry in IPBWT that does not compare less than ⟨*Y, l*⟩ . We are free to choose any implementation for how to find that lower-bound location; for example, since the IPBWT is sorted, we could do a binary search over the entries of the IPBWT. Fig. 9a shows how to create an IPBWT with *K* = 3.

Given *f* ^*K*^, we can define Supplementary Alg. 2 to perform *exact search* using IPBWT. For example, using the reference sequence and IPBWT from Fig. 9a, let the query sequence be ATTA. We split this into two chunks: ATT and A (line 3). We first use the RMI to find the lower bound locations of (*A*$*A*, 0) and (*ATT*, |*R*|), which are 1 and 5, respectively. Subsequently, we use the RMI to find the lower bound locations of (*ATT*, 1 )and (*ATT*, 5), which are 3 and 5. Our algorithm gives the interval [3, 5) . We can confirm that ATTA can be found in position 3 and 4 of the BW-Matrix.

**Figure 9:**
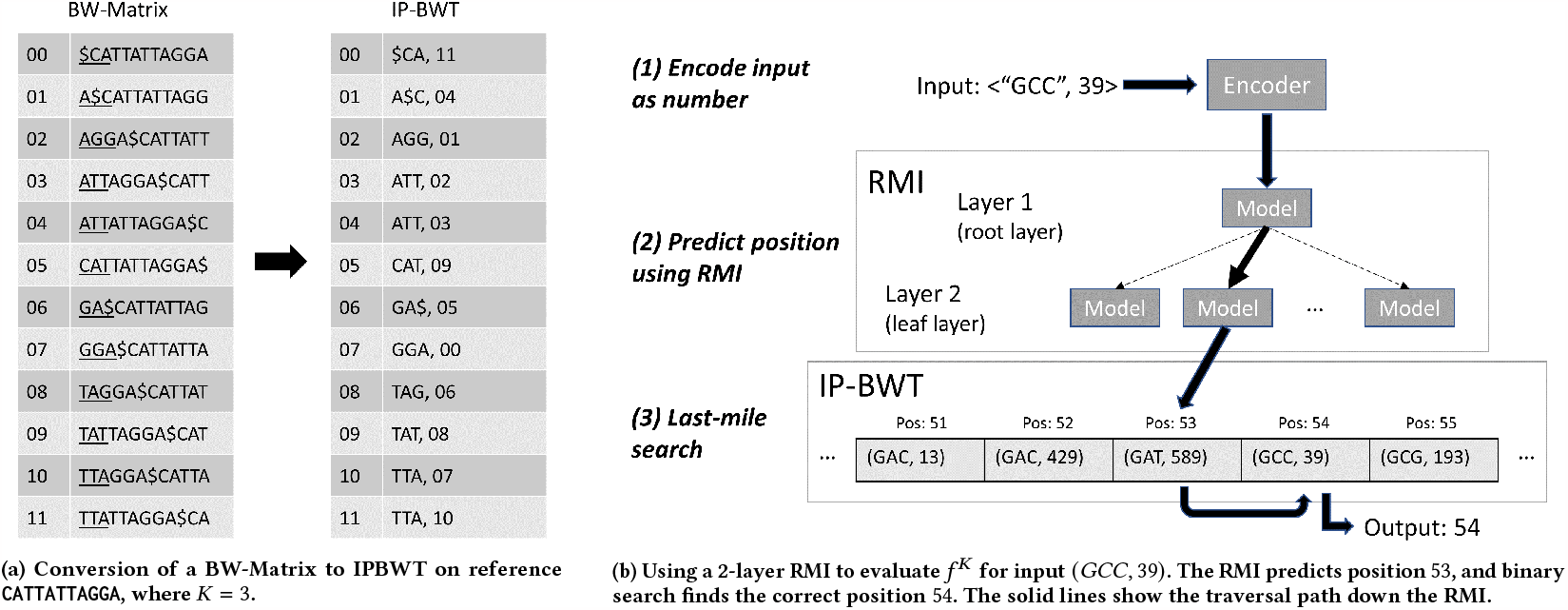
Components of the LISA index for the *exact search* algorithm

#### 5.6.2 Faster Chunk Processing using RMI

Using the IPBWT, we are able to process the query sequence in chunks of *K* bases at a time. However, when processing each chunk, we must evaluate the function *f* ^*K*^ . Using a binary search over the IPBWT takes *O* (log |*R*|) time. Therefore, the overall runtime of *exact search* using IPBWT with binary search for *f* ^*K*^ is 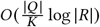 . For large reference sequences, this might be slower than *backward-search* using the FM-index. Therefore, we use a learned approach to accelerate *f* ^*K*^ . In particular, *f* ^*K*^ is a model that maps an input key –⟨ *Y, l*) ⟩ – to its position in the sorted IPBWT. Therefore, it can be replaced by a machine learning model. Based on this insight, in LISA, we use knowledge of the distribution of subsequences within the reference sequence to create a learned index structure to model *f* ^*K*^ . We experimented with three of the prominent learned index structures – Recursive Model index (RMI) [20, 39], Piecewise Geometric Model Index (PGM) [36, 40], and Radix Spline index (RS) [41]. We found that a 2-layer RMI worked the best for our application, which is in line with the recently published benchmarking study comparing various learned indexes [42, 43].

Therefore, we model *f* ^*K*^ using an RMI, which is a hierarchy of models that are quick to evaluate [20]; the RMI conceptually resembles a hierarchical mixture of experts [44]. A 2-layer RMI – that only has root and leaf layers – has a single model at the root layer. For any input key, the model at root layer is used to predict the correct model to use at the leaf layer. The predicted model at the leaf layer is used to predict position of the key in a sorted array and a range of positions in which the key is guaranteed to occur. If the key is not found in the predicted position, a last mile search is conducted in the provided range to find the key. Fig. 9b illustrates, with an example, how we use a 2-layer RMI to implement *f* ^*K*^ in three steps:

1. Since the RMI only accepts numbers as keys, we first convert the input ⟨*Y, l*⟩ into a number. Since there are only 4 bases, each base can be represented using 2 bits. Therefore, we convert the DNA sequence *Y* into an integer, *I* (*Y*), with 2*K* bits by concatenating the bits of the individual bases together and creating the key as 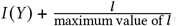 which is a double precision (64 bit) floating point number.
2. We give the encoded key to the RMI and traverse down the layers of the RMI to a leaf model. The leaf model predicts the position in the IPBWT where it expects to find the input key and a range in which we are guaranteed to find the input key.
3. If the predicted position does not contain the input pair, we use binary search over the range to find the actual position of the key.

Note that this learning-based approach to modeling *f* ^*K*^ guarantees correctness; LISA will produce exactly the same results as using backward search with FM-index. Instead of a floating point key, we can also create an integer representation as key by just concatenating the bits of *l* to *I* (*Y*), but that may result in keys of length greater than 64 bits and will need more than one step to compare – comparing 64 bits in each step. Another option is to use the most significant 64 bits of the integer representation as key, but that results in losing differentiating information when the most significant 64 bits are zero. Therefore, we represent the key as a double-precision (64 bits) floating point number and train the model over an array where each entry is a double-precision representation of the corresponding entry in the IPBWT. However, we perform last-mile search over the original IPBWT so as to ensure correctness. Note that we have a special case for handling the sentinel letter $ while maintaining this 2-bit encoding. Figure 7 describes LISA’s chunking of *k* bases resulting in fewer backward search iterations. Figure 8 describes an example of 4 *k* bases backward extend steps and corresponding change in SA interval.

**Figure 7:**
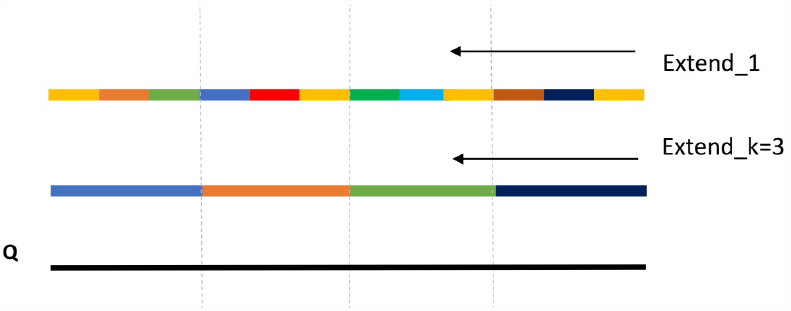
Comparison between LISA and FM-index based query (*Q*) search: extending query match *k* bases at-a-time using LISA vs 1 base at-a-time using FM-index.

**Figure 8:**
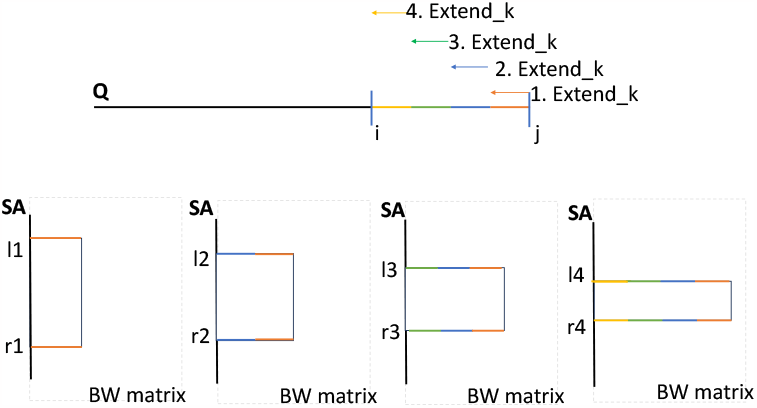
Example 4 steps of extending query (*Q*) match using LISA *k* bases at a time and the corresponding change in SA interval.

#### 5.6.3 Parameter Selection

The number of leaf nodes is an important parameter for the RMI, as shown in Supplementary Table 6. Larger number of leaf nodes results in better accuracy and smaller ranges for last mile search – thereby, smaller number of iterations of the binary search and better throughput – at the cost of larger memory consumption. Another important parameter is *K*, as shown in Supplementary Table 7. For this experiment, we used a seed length that is divisible by the values of *K* to ensure that no length is at a disadvantage. Increasing *K* results in smaller number of steps of binary search, but would consume more memory. Moreover, 64 bits may not be sufficient to represent the corresponding keys for very large values of *K*, which might lead to information loss and worse model accuracy resulting in longer ranges to perform binary search in.

#### 5.6.4 Performance Comparison with TAL using Synthetic Queries

In case of real datasets, few queries may not find a match in the reference. In those cases, we may not need to process the entire query. The FM-index based approach (Supplementary Alg. 1) can stop as soon as an additional base results in no matches, while the LISA-based approach can stop only after a chunk is processed and returns no matches, thereby, processing more bases than the FM-index based approach. While LISA gets significant performance gain over FM-index based approach for real queries as shown in Sec. 2, it can achieve even higher speedup if all the queries have a match. Therefore, to showcase the maximum benefit of LISA, here we perform experiments using synthetic queries directly extracted from the corresponding reference sequence. We use queries of length 22, as that is the default length of seeds for Bowtie2. Supplementary Fig. 2 shows that LISA achieves 1.8 −2× and 1.9 −2× speedup over TAL on a single thread and single CPU socket, respectively – therefore, on an average, achieving higher speedup compared to real queries.

### 5.7 The SMEM Search Problem

*SMEM search* finds all the SMEMs between a query and a reference sequence as defined in the following. A *maximal exact match* (MEM) is an exact match between substrings of two sequences that cannot be further extended in either direction. An SMEM is a MEM that is not contained in any other MEMs on the query sequence [45]. Thus, SMEM search produces exact matches of variable lengths by definition. Figure 10 shows an example SMEM matches between a query *Q* and reference sequence *R*. For reference sequence *R* and query sequence *Q*, we say *Q* [*i, j* ] is an SMEM exactly when *Q* [*i, j* ] has one or more matches in *R* but ∀ *i* ^′^, *j* ^′^ such that *i* ^′^ *< i, j* ^′^ *> j*, it is the case that *Q* [*i* ^′^, *j*], *Q* [*i, j* ^′^ ] and *Q* [*i* ^′^, *j* ^′^] have no matches. Therefore, no two SMEMs can share the same upper-bound or lower-bound.

**Figure 10:**
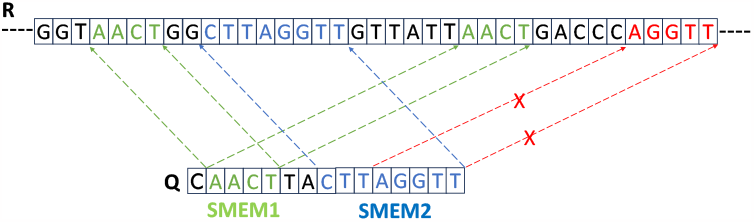
SMEMs between query and reference sequence: For query *Q*, SMEM1 matches to 2 places, and SMEM2 matches to 1 position in reference sequence *R*. The Red arrows point to a matching substring (a MEM) between *Q* and *R*, but it does not qualify to be a SMEM as it is subsumed by a bigger match (SMEM2).

Typically, |*R*| ≈10^9^ bases; e.g., the length of the human genome is nearly 3 × 10^9^. On the other hand, |*Q*| is typically a few hundred bases (*<* 300 bases); e.g., a common query length in BWA-MEM for *SMEM search* is 151. To cover both the strands of the DNA, we look for matches of |*Q* | in both *R* and the reverse complement of *R*.

### 5.8 Prior work related to SMEM Search

The algorithm for *SMEM Search* [6, 45] in BWA-MEM uses a bidirectional FM-index in which *R* is concatenated with the reverse complement of *R* and the FM-index of the concatenated sequence is built. The algorithm performs the following steps.

1. It begins at the end of the query, backward extending by one letter at a time until there is at least one exact match. For backward extension, it uses FM-index employing an algorithm similar to Supplementary Alg. 1. At the end of backward extension, it returns the corresponding match found – as an SMEM.
2. Then it starts building the next SMEM from the position right next to the SMEM found. First, it forward extends using the benefit of bidirectional FM-index to reduce the problem of forward extension to that of backward extension. It performs forward extension until there is at least one match, while maintaining all the intermediate forward extensions. For each of the intermediate forward extensions, it performs backward extension and outputs the longest match as SMEM.
3. It performs step 2 repeatedly until the entire query is processed.

Due to step 2, the algorithm has *O* (|*Q*| ^2^) time complexity.

The search of Maximal Exact Matches (MEM), a closely related problem, has been well studied in literature using suffix arrays, sparse suffix arrays, FM-indexes, k-mer indexes, bloom filters, etc. [46– 53]. In particular, [46] proposes the use of enhanced suffix arrays [54], a combination of FM-index and lcp-interval tree, to search for MEMs. For our LISA based SMEM algorithm – we (1) adapt the algorithm proposed in [46] to the SMEM search problem to develop an *O* (|*Q*|) algorithm to search for SMEMs – to the best of our knowledge, no prior work has proposed an *O* (|*Q*|) algorithm for SMEM search, (2) apply the learned approach and (3) develop an efficient hardware-aware implementation – to achieve significant performance gains.

### 5.9 SMEM-LISA: SMEM Search based on LISA

In this section, we first introduce a few properties of SMEM key to our algorithmic discussion. Subsequently, we briefly outline our new algorithm along with the high-level idea behind it. Finally, we give a detailed description of our algorithm. We define our LISA index used for SMEM search as the tuple, *L* = ⟨*F, I PBW T, RMI, T, LCP*⟩, where *F, I PBW T*, and *RMI* are as defined above and *T* and *LCP* will be defined in this section.

#### 5.9.1 Monotonicity and Consistency Properties of SMEMs

Compared to the FM-index based SMEM algorithm, where SMEMs are found by extending candidate substrings forward and backward, our new algorithm extends as well as ‘shrinks’ substrings to allow for efficient SMEM traversal. Before we describe the algorithm, we present a few key observations about the underlying structure of SMEMs:

##### Definition

*For query sequence Q, we denote a substring of Q, Q* [*i, j* ], *and its SA interval*, [*l, r* ), *as a tuple* ⟨*i, j, l, r* ⟩.

##### Lemma

(The monotonicity property of SMEM). *For any two different SMEMs Q* [*i*_1_, *j*_1_] *and Q* [*i*_2_, *j*_2_], *we have i*_1_ *< i*_2_ ⇔ *j*_1_ *< j*_2_.

Proof. Without loss of generality, say *i*_1_ *< i*_2_. If *j*_1_ ≥ *j*_2_, then *Q* [*i*_2_, *j*_2_] is contained in *Q* [*i*_1_, *j*_1_] and thus, can not be an SMEM. □

From this, we know that SMEMs appear in ‘strictly increasing’ order; *i* increases exactly if *j* increases. Therefore we can order them with either *i* or *j* (both are equivalent), and define the notion of adjacency.

##### Definition

*We say two different* SMEM*s Q* [*i*_1_, *j*_1_], *Q* [*i*_2_, *j*_2_] *are* ***adjacent*** *if there doesn’t exist another* SMEM *Q* [*i*_3_, *j*_3_] *such that i*_3_ *is between i*_1_ *and i*_2_. *If i*_1_ *< i*_2_, *we say Q* [*i*_1_, *j*_1_] *is the* ***left-adjacent*** SMEM *of Q* [*i*_2_, *j*_2_], *and Q* [*i*_2_, *j*_2_] *is the* ***right-adjacent*** SMEM *of Q* [*i*_1_, *j*_1_].

##### Lemma

(The consistency property of SMEM). *Consider any two adjacent* SMEM*s Q* [*i*_1_, *j*_1_] *and Q* [*i*_2_, *j*_2_], *where i*_1_ *< i*_2_. *For all j* ∈ [ *j*_1_ + 1, *j*_2_], *we have that Q* [*i*_2_ − 1, *j* ] *has no match in R*.

Proof. If *Q* [*i*_2_ − 1, *j* ] has a match in *R*, then there exists some SMEM *Q* [*i*^∗^, *j* ^∗^] s.t. *i*^∗^ ≤ *i*_2_ − 1 and *j* ^∗^ ≥ *j* . By the monotonicity property, *i*^∗^ ≤ *i*_2_ − 1 ⇒ *j* ^∗^ *< j*_2_. Also, because *Q* [*i*_1_, *j*_1_] is left-adjacent to *Q* [*i*_2_, *j*_2_], we have *j* ^∗^ *< j*_2_ ⇒ *j* ^∗^ ≤ *j*_1_. But *j* ^∗^ ≥ *j* ≥ *j*_1_ + 1.

The Consistency Property is the most crucial observation to our new algorithm. It says that if we begin with *Q* [*i* = *i*_2_, *j* = ] *j*_2_ and start decrementing *j*, the lower-bound *i*_2_ will remain the lowest possible bound to form a match, until we reach *j* = *j*_1_. This suggests an efficient way to traverse adjacent SMEMs. Say we are at *Q* [*i* = *i*_2_, *j* = *j*_2_] and want to move to its left-adjacent SMEM *Q* [*i*_1_, *j*_1_]. What we can do is keep decreasing *j* while testing if *Q* [*i* − 1, *j* ] still has no match in *R*. The first *j* yielding one or more matches for *Q* [*i* − 1, *j* ] in *R* must be *j*_1_. Thus we have reached *Q* [*i* = *i*_2_, *j* = *j*_1_]. Then we can simply apply backward extension to reach *Q* [*i* = *i*_1_, *j* = *j*_1_].

#### 5.9.2 The New *SMEM Algorithm*

Figure 11 pictorially depicts the extend and shrink phases of LISA based SMEM search algorithm. Supplementary Alg. 3 utilizes the Consistency Property to find SMEMs of a query. Say *Q* has *n* SMEMs, in increasing order of *i*: *Q* [*i*_1_, *j*_1_], *Q* [*i*_2_, *j*_2_], …, *Q* [*i*_*n*_, *j*_*n*_], where *i*_1_ = 0 and *j*_*n*_ = |*Q*| −1. We start from *i* = |*Q*| and *j* = *j*_*n*_ = |*Q*| −1 (line #3). We start with an extend phase (lines #4-5) and then alternate between a shrink phase (line #8) and an extend phase (line #9) in a while loop (lines #7-10). Each extend phase, given *Q* [*i*_*p*_, *j*_*p* − 1_], finds us *Q* [*i*_*p* − 1_, *j*_*p*− 1_], and shrink phase, given *Q* [*i*_*p*_, *j*_*p*_ ], finds us *Q* [*i*_*p*_, *j*_*p*− 1_ ]. Therefore, the while loop traverses all SMEMs in reverse order.

**Figure 11:**
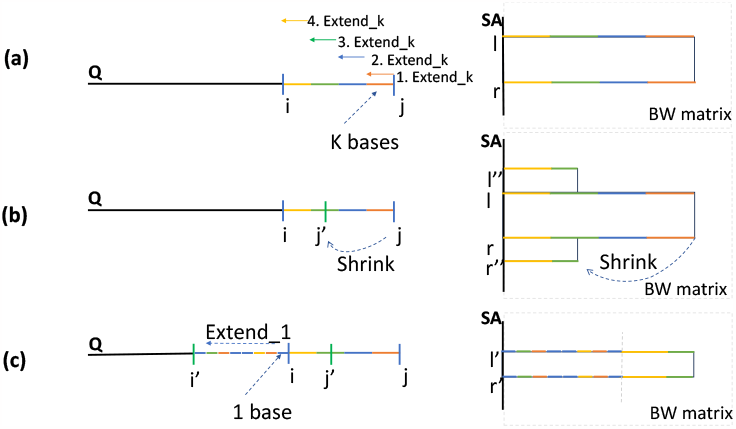
Example showcasing Extend and Shrink phases of Supplementary Algorithm 3. The figure depicts query *Q*, SA interval and the length of match in the SA interval. (a) first SMEM ⟨*i, j, l, r* ⟩of query *Q* resulting from four backward extends of *k* bases each, (b) the following shrink phase resulting in partial match ⟨*i, j* ^′^, *l* ^′′^, *r* ^′′^⟩, (c) extends the partial match 1 base at a time resulting in the second SMEM ⟨*i* ^′^, *j* ^′^, *l* ^′^, *r* ^′^⟩.

#### 5.9.3 Extend Phase

The task of the extend phase is fairly straight-forward. Since the upper-bound *j* of the SMEM is known, we can simply extend leftward until right before no match. The first extend phase may be long as we are starting from scratch and need to extend until we find one SMEM. All subsequent extensions are done after a shrink phase and start with a partial match already and thus, may not need long extensions. Based on this insight, in the first extend phase, we use the LISA index to extend in chunks of *K* bases at a time until we can not extend by *K* bases anymore (line #4). Subsequently, we extend by one base at a time until there are matches (line #5). For extensions after shrink phases, we only use the FM-index based one base at a time method (line #9). Supplementary Alg. 4 and Supplementary Alg. 5 detail how we extend one base at a time using FM-index and *K* bases at a time using LISA, respectively. These are similar to Supplementary Alg. 1 and Supplementary Alg. 2, respectively, but instead of returning *ϕ* when they can’t match the entire query, they return the positions up to which they could find matches.

#### 5.9.4 Shrink Phase

Given an SMEM, *Q* [*i, j*], according to the Consistency Property, we need to find the largest index *j* ^′^ *< j* such that *Q* [*i, j* ^′^ ] is left extendable (that is, *Q* [*i* −1, *j* ^′^] has a match in *R*) and the suffix interval of *Q* [*i, j* ^′^] . This requires us to support two functionalities:

- For any *Q* [*i, j* ], find if it is extendable to the left.
- Given *Q* [*i, j* ] and its SA interval [*l, r* ), find the SA interval of *Q* [*i, j*^′^ ].

Lemma (The Extendability Property). *For a string Q i, j with SA interval [l, r*), *if we keep removing bases from the right, its left extendability remains the same if the SA interval remains the same. In other words, for any j* ^′^, *such that Q* [*i, j* ^′^] *has the same SA interval as Q* [*i, j*[, *Q*[*i* −1, *j* ^′^] *has a match in R if and only if Q*[ *i* −1, *j*] *has a match in R*.

Proof. Without loss of generality, if *j* ^′^ *< j* and *Q* [*i, j* ] and *Q* [*i, j* ^′^] have the same SA interval, that means *Q* [*i, j* ^′^] is always followed by *Q* [ *j* ^′^ + 1, *j* ] in *R*. So, if *Q* [*i* − 1, *j* ] has a match in *R, Q* [*i* − 1, *j*^′^ ] also has a match and if *Q* [*i* − 1, *j* ] does not have a match in *R, Q* [*i* − 1, *j* ^′^] also does not have a match. □

Therefore, the extendability property only depends on the SA interval [*l, r*) and the base *Q* [*i* −1] . The corollary of the extendability property is that for shrink phase, we need to remove bases from the right of *Q* [*i, j*] until the SA interval changes and then check if the resultant string is left extendable. Our solution to this problem is based on Enhanced Suffix Arrays (ESA) [54] as proposed in Section 4 of [46].

##### Enhanced Suffix Arrays

Fig. 12 shows the ESA consisting of the BW-Matrix, LCP array, suffix tree and lcp-interval tree of our example reference sequence. The LCP (Longest Common Prefix) array, *LCP*, is an array of size |*R*|+ 1 consisting of integers in the range 0, *R* . *LCP i* is defined as the length of the longest common prefix of BW-Matrix entries at *i* and *i* −1; *LCP* [0] and *LCP* [|*R*|] are set to 0 as there are no BW-Matrix entries for positions 1 and |*R*| [54]. A suffix tree [55] of a DNA sequence is a compressed trie containing all the suffixes of the sequence as keys and positions in the sequence as their values. Each node in the suffix tree represents a substring of the reference sequence, formed by concatenating the labels of the edges from the root node up to that node. Each leaf node represents a suffix of |*R*| . Therefore, number of leaf nodes is equal to |*R*| . An lcp-interval tree can be created by replacing each node label of a suffix tree with the SA interval of the corresponding substring it represents. For all possible strings with non-zero matches, its SA interval can only be one of the 19 possibilities corresponding to the 19 nodes in Fig. 12c. Moreover, these 19 intervals form a certain hierarchical tree, where an edge from *x* to *y* means that *y* is the first non-*x* SA interval we get if we keep removing bases on the right. Each non-leaf node is exactly a union of its children nodes, and the intervals of these children nodes are disjoint. Given *Q* [*i, j* ] and its SA interval [*l, r* ), the SA interval of *Q* [*i, j* − 1] could either stay as [*l, r* ) or it could be the SA interval corresponding to the parent node of the node with label [*l, r* ).

**Figure 12:**
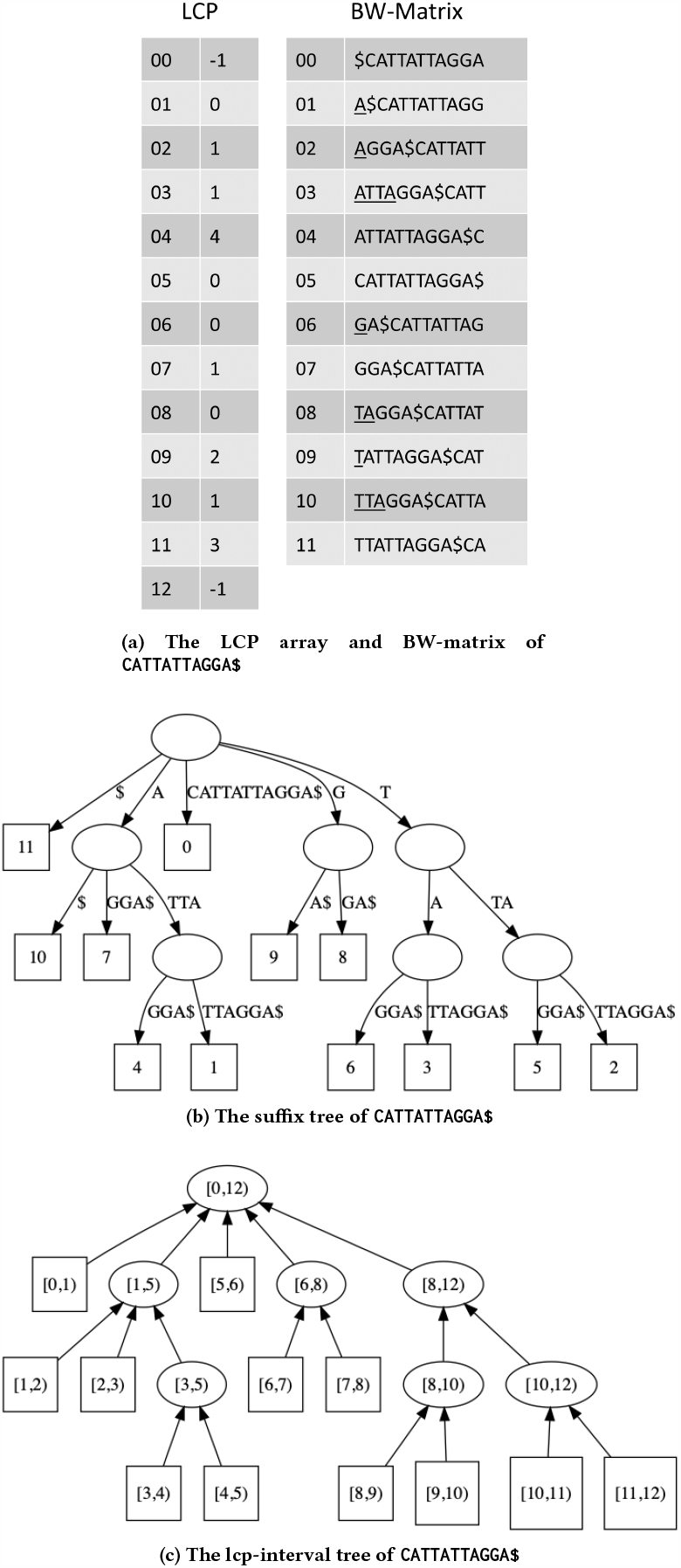
The Enhanced Suffix Array, Suffix Tree, and Interval Tree of *R* ← CATTATTAGGA$.

We make the following enhancements to the design of the lcp-interval tree for its efficient representation and traversal.

- **Binary lcp-interval tree :**To store the lcp-interval tree, we need to store the parent node at each of the node requiring significant additional memory. Instead, we opt to insert a few “dummy” nodes to make it binary while maintaining the properties of the lcp-interval tree (Fig. 13). It is easy to see that there are now exactly |*R*| − 1 non-leaf nodes and |*R*| leaves. To store all non-leaf nodes of the binary tree, we build a size-(|*R*| −1) array, called *T*, and each non-leaf node is stored at the position where the intervals of its two children meet. For example, the node [1, 5) will be stored at position 2. This is equivalent to storing these nodes in the order of the in-order traversal. This also means that for each non-leaf node stored at position *p, LCP* [*p*] is the length of the string represented by the node. For example, for the node corresponding to the interval [10, 12], its children [10, 11) and [11, 12) meet at 11. So, [10, 12) is stored at position *p* = 11. *LCP* [11] is 3 and from Figures 12b and 12c, we can see that [10, 12) represents the string *TT A* of length Later, we will show that we don’t need to store the leaves.

**Figure 13:**
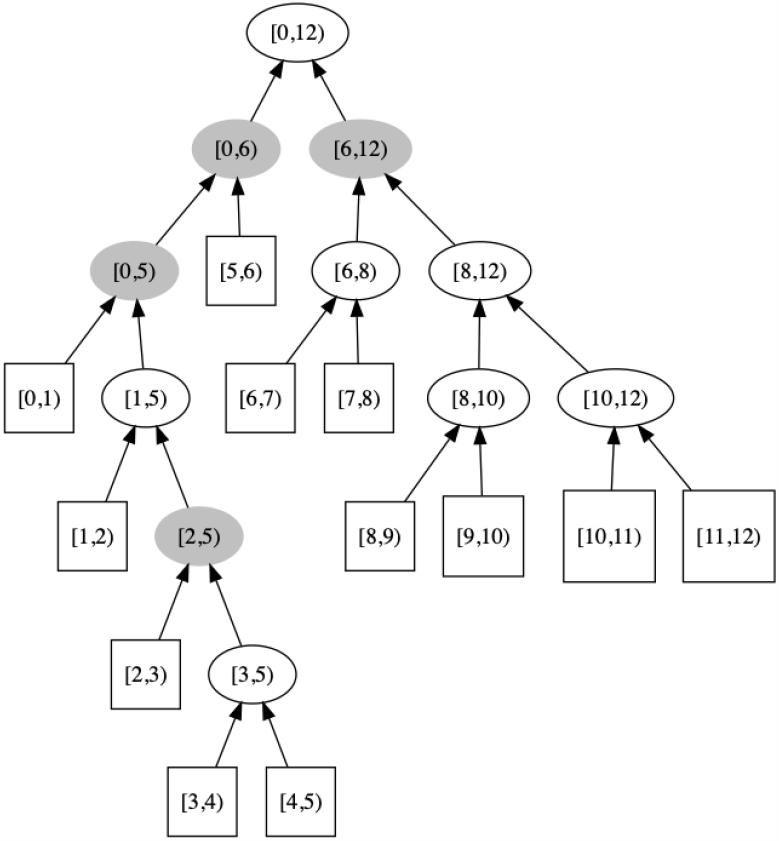
The binary lcp-interval tree of CATTATTAGGA. The dummy nodes are colored gray.
- **Storing interval size instead of interval**. In the array *T*, for each non-leaf node, we don’t need to store the interval. Instead to save memory, we store only the size of the node’s interval. The *p*-th such entry is denoted as*T* [*p*] .*size*. The interval can be reconstructed from size and *p*. This size alone is sufficient to climb up the tree (Supplementary Alg. 6). Suppose we are at some node [*l, r* ). We know that its parent must be stored at either index *l* or index *r* . Moreover, the index out of *l* and *r* not storing the parent stores some further ancestor, whose interval size must be larger than that of the parent. Using this, to find the parent of [*l, r*), we simply compare *T*[ *l*] .*size* and *T* [*r*] .*size*. If the former is smaller, then the parent node is stored at index *l*, so the parent’s interval is [*r*− *T* [*l*] .*size, r*) ; otherwise the parent node is stored at index *r*, so its interval is [*l, l* +*T* [*r*] .*size*) . Therefore, Supplementary Alg. 6 allows us to remove the minimum number of bases such that the SA interval changes and also provides the corresponding SA interval. In order to quickly check if the resultant shrunk string is left extendable, for each entry in *T*, we store a 4 bit entry called *ext* to store whether it is left extendable by *A, C, G* and *T* . For the dummy nodes, these 4 bits are all set to false. Now, given SMEM *Q* [*i, j*] and its SA interval [*l, r*), the objective of the shrink phase becomes finding the nearest ancestor of [*l, r*) in the binary interval tree such that the *ext* bit for *Q* [*i*− 1] is set to true.
- **Not storing leaves**. Finally, we note that the leaves are not stored. This is because the SMEM *Q* [*i, j*] is not left extendable by definition. So its SA interval must always climb up at least one edge to reach the shrink phase’s answer, and thus leaves are never the answer.

##### Implementing Shrink Phase Using Shrink Operation

Our algorithm for shrink phase is presented in Supplementary Alg. 7. We start by one shrink operation (line #2) and while the *ext* bit corresponding to *Q*[ *i* − 1] is still false, we keep performing shrink operations (lines #3-4). Since *LCP*[ *p*] is the length of the string represented by the node at position *p*, we can get *j* ^′^ from *i* and *LCP* [*p*] (line #5).

### 5.10 Performance Evaluation of SMEM Acceleration

Supplementary Section A.5 shows that, in practice, the time complexity of our LISA based SMEM algorithm is sub linear in |*Q*| . In this section, we evaluate the performance benefits.

#### 5.10.1 Quantifying the benefits of ESA and learned approach

In these experiments, we compare TAL with two versions of our implementation: (1) Linear: the ESA based linear algorithm with the learned approach based enhancement turned off and 2) SMEM-LISA: ESA based linear algorithm enhanced with learned approach. For a fair comparison with the implementation in TAL that is well tuned for the underlying CPU, we developed hardware-efficient implementations for Linear and SMEM-LISA and use those for comparison. Supplementary Fig. 3 shows that our linear algorithm achieves significantly high speedups of 3.9 5.2 over TAL. The learned approach achieves additional performance gain, achieving a total speedup of 5.7 − 7.5×.

#### 5.10.2 Performance Comparison with TAL using Synthetic Reads

In these experiments, we compare our performance with TAL with respect to variation in read error rates by generating synthetic reads of various error rates ranging between 1 − 10% using wgsim [56]. Supplementary Fig. 4 shows that we achieve up to 9.6× and 6× speedups over TAL on a single thread and a of using enhanced suffix arrays, learned index based approach and an efficient hardware aware implementation. While we achieve significant speedups even with high error rates, the speedup is higher when the error rate is lower. This is because LISA is used to extend *K* bases at a time only for the first extension. The smaller the error rate, the higher the chances of the first extension to be longer and LISA to be used for a larger portion of the read. Note that, for the latest short-read sequencers, a majority of bases have error rates of just 0.1%. This is reflected in higher speedup achieved by our approach on real datasets (Sec. 2).

### 5.11 Bringing it all together: Acceleration of Seeding Phase of BWA-MEM2

#### 5.11.1 Background on BWA-MEM2

BWA-MEM is a highly popular genomics software tool to map short DNA sequences to a reference sequence, with more than a million downloads/installs. BWA-MEM2 is an accelerated version of BWA-MEM that uses the underlying CPU architecture efficiently. BWA-MEM2 is a drop in replacement of BWA-MEM with identical output and up to 3x faster performance [7]. BWA-MEM2 runtime is dominated by three major phases: 1. Seeding, 2. Chaining, and 3. Sequence alignment.

#### 5.11.2 The Seeding Phase

The seeding phase involves searching for the exactly matching substrings between a read and a reference sequence and consumes 30 − 40% of the BWA-MEM2 runtime. The seeding phase of BWA-MEM2 consists of three kernels (say K1, K2, and K3) each of which is a variant of *SMEM or exact search* algorithms. Supplementary Alg. 14 presents LISA-based seeding phase in BWA-MEM2. Next, we discuss the FM index-based implementations of the three kernels in BWA-MEM2 and present their accelerated LISA-based implementations.

#### 5.11.3 SMEM Search (K1)

In BWA-MEM2, the first seeding kernel finds all SMEMs of a query *Q* with additional constraints on the minimum seed length *min*_*seed*_*len*. It uses FMI-based SMEM search algorithm, that is described in Section 5.8. Recall that the FMI-based algorithm processes one-base at a time and has *O* (|*Q*| ^2^ ) time complexity. The default value of *min*_*seed*_*len* is 19 which translates to - a search of all possible SMEMs of length ≥ 19.

The LISA-based algorithm for K1 uses a parameterized variant (Supplementary Alg. 8) of the LISA-based SMEM algorithm described earlier (Supplementary Alg. 3). This variant uses the parameter, minimum seed length *min*_*seed*_*len*, as input and ensures that the SMEMs found are longer than *min*_*seed*_*len* (lines #6 and #13). There are other parameters to this variant that we discuss below.

#### 5.11.4 Re-Seeding (K 2)

For each SMEM *SK* 1 : ⟨*i, j, l, r*⟩ found in K1 where the seed length (*j* −*i* +1) *> re*_*seed*_*thresh* (the re-seeding threshold), the reseeding kernel *K* 2 finds another set of SMEMs that overlap with the middle base of *SK* 1 and has more matching frequency than *SK* 1 . Specifically, *K* 2 adds an *SMEM SK* 2 : ⟨*i* ^′^, *j* ^′^, *l* ^′^, *r* ^′^⟩ to the output if *i* ^′^≤ *m* ≤*j* ^′^ where *m* = (*i* +*j*) /2 is the mid point of *SK* 1 (*pivot* ) and *j* ^′^ −*i* ^′^ +1 *> min*_*seed*_*len* and *r* ^′^ −*l* ^′^ *> r* −*l* . The default value of *re*_*seed*_*thresh* is 28. While the matches output by K2 are not technically SMEMs, in BWA-MEM2, K2 uses the same SMEM search algorithm as K1 by parameterizing with the above mentioned thresholds. In BWA-MEM2, K2 starts the search from the pivot point using bi-directional FMI.

However, LISA-based SMEM search does not directly support SMEM search from any arbitrary position – it can only start from the end of a query and progresses the search towards left. A trivial and inefficient LISA-based solution would be to start SMEM search from the end of the query, collect all possible *SMEM*s and then filterout the ones crossing the *pivot* point. This is compute intensive and redundant. Instead, we start from the *pivot* and find the right-most position *j* ^′^ using forward FM-index based search (same as backward search with reverse-complemented query) such that *Q* [*pivot, j* ^′^] is a match in *R* and *Q* [*pivot, j* ^′ +^1] is not a match (Supplementary Alg. 14, line #7). In other words, *j* ^′^ is the right-most position among all SMEMs to be found in K2 and it is guaranteed that no *SMEM* is to be found beyond *j* ^′^. From this right-most position, we use Supplementary Alg. 8 with *pivot* and the frequency threshold, and abort the search once all *SMEM* crossing the *pivot* are found (line #10). Supplementary Alg. 9 and 10 (line 3) track the frequency of the matching string using the size of suffix array interval and stops the search when the frequency drops below *r* − *l* .

#### 5.11.5 Seed Strategy (K 3)

The seed-strategy kernel of BWA-MEM2 finds yet another set of non-overlapping matches with *minimum seed length min*_*seed*_*len* and with the number of occurrences under max-hit threshold*max*_*hit*_*thresh. min*_*seed*_*len* and*max*_*hit*_*thresh* are both set to 20 by default. The seed strategy works as follows: Given a query *Q*, the minimum seed length *min*_*seed*_*len* and the max-hit threshold *max*_*hit* _*thresh*, the kernel first tries to find exact matches of *Q* [0, *min*_*seed*_*len*− 1] . Suppose *n*_*match*_ denotes the number of matches of *Q* [0, *min*_*seed*_*len* −1] . There are three possible outcomes of the search, and the seed strategy decides the next step accordingly:

1. *n*_*match*_ ≥*max*_*hit* _*thresh*: Continue the search by appending one base at a time till *n*_*match*_ ≥ *max*_*hit* _*thresh*. If *i* is the first position such that *i, Q* [0, *i*] satisfies *n*_*match*_ *< max*_*hit* _*thresh*, add *Q* [0, *i*] to the output and start a fresh search for *Q* [*i* +1, *i* +*min*_*seed*_*len*] .
2. 0 *< n*_*match*_ *< max*_*hit* _*thresh*: The search is successful, add *Q* [0, *min*_*seed*_*len* − 1] to the output and start a fresh search for *Q* [*min*_*seed*_*len*, 2 × *min*_*seed*_*len* − 1].
3. *n*_*match*_ ≤ 0: The search for *Q* [0, *min*_*seed*_*len* − 1] is unsuccessful. Start a fresh search for *Q* [*min*_*seed*_*len*, 2 × *min*_*seed*_*len* − 1].

In seed strategy, BWA-MEM2 performs FMI based search through forward extension and appends one base at a time in the rightward direction. It is important to note that, forward-extend using bi-directional FMI search has to maintain SA intervals on both forward and reverse strand of the reference sequence and therefore it is computationally costlier than the traditional backward-extend FMI search. Supplementary Alg. 13 presents FMI-based forward-extend step processing one base at a time. Given a query *Q*, the tuple ⟨*i, j, l*_1_, *l*_2_, *s*⟩ represents that the subquery *Q* [*i, j*] is a match in the reference sequence with l1 and l2 are the lower bounds of SA intervals on the forward and the reverse strand, respectively, and *s* is the size of the matching intervals. The function *f* ^′^ defines one step of bi-directional FMI search in forward direction. *f* ^′^ receives two arguments: 1) the complement of a base to extend, i.e., 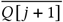, and 2) a tuple with SA intervals on both the strands, ⟨*l* 1, *l*2, *s*⟩ and returns a new set of SA intervals matching *Q* [*i, j* +1] . Compared to FMI search function *f*, the new function *f* ^′^ requires more operations to obtain the SA intervals on the both strands.

Supplementary Alg. 11 presents our LISA-based seed strategy for K3. We train LISA index with *K* = *min*_*seed*_*len*, i.e., the minimum seed length of K3, and we perform exact-search for the first *min*_*seed*_*len* bases in one-shot using RMI. If the search is successful, in majority of the cases the K-sized matches also satisfy the max-hit threshold criteria and the match is added to the output (line 5-8). Occasionally, if the maximum frequency threshold is still not met, the search continues with forward-extend by one base at a time using FM-index (line 10). Note that, forward-extend with FMI requires SA intervals of both strands. So LISA ensures that the SA intervals are obtained for both strands (if needed) by performing one more LISA search with the reverse-complemented chunk of size K.

### 5.12 Hardware-aware Design and Implementation

In this section, we describe our design for efficient utilization of the underlying CPU hardware so as to achieve maximum throughput for our LISA-based algorithms.

#### 5.12.1 Motivation for using Software Prefetching

A common pattern across all the algorithms discussed in this paper is that they all perform frequent memory accesses over large data structures (that cannot fit in cache) and very little compute, making them memory access bound. CPUs have hardware prefetchers that can prefetch data from memory in advance if there is a predictable pattern of memory addresses. However, for these algorithms, the addresses of the memory accesses are pseudo-random — i.e.; there is no clear pattern across subsequent accesses — and computation in one step decides the memory addresses for the next step, making it impossible for the hardware prefetchers to prefetch them. For example, in Supplementary Alg. 1, in each iteration of the loop, two positions of *O* are accessed and the computation decides which two positions would be accessed in the next iteration. The positions of *O* accessed in subsequent iterations can be vastly different from each other. Similarly, for the computation of *f* ^*K*^ shown in Fig. 9b, the computation of the level 1 of the RMI reveals which model of level 2 has to be used and the parameters of that model have to be read. In the last mile search, we use binary search, where each iteration reveals which half of the search space would be processed next.

Therefore, for every pseudo random memory access, we have to pay the cost of getting data from memory (memory latency) – making the algorithms memory latency bound. This necessitates the use of software prefetching — software instructions used to specify which memory addresses should be prefetched to cache in advance. These instructions are non-blocking; i.e.; they start the process of prefetching and return the control back so that other instructions can be run in the meantime, thus, allowing for overlap of memory access with computation. Each step decides the memory addresses for the next step and the little compute that is there in each step can not hide the memory latency for the prefetch of the next step. Therefore, we use the design described in [32] for backward search algorithm for *exact search* (Supplementary Alg. 1) and apply it to the Exact-LISA (Supplementary Alg. 2) and SMEM-LISA (Supplementary Alg. 3) algorithms. Compared to backward search algorithm, LISA based algorithms are a lot more complex with many varied steps each requiring a different number of memory accesses making it particularly challenging.

#### 5.12.2 Our Design for Software Prefetching

Here, we describe a generic design (Figure 14) that we have applied to each LISA-based algorithm. There is no data dependency across queries and the queries can be processed in parallel. On the other hand, processing of a query consists of several steps, which need to be executed in a sequential manner because of data dependency across steps. We start by formally defining a step. A step is a unit of work that receives a set of memory addresses as input, reads data from the memory addresses, performs compute on the data and generates a new set of memory addresses for the next step. The first and the last step are a little different. The first step (setup step) does not receive addresses to access data from memory and the last step produces the query output instead of memory addresses. Important to note that no step should access a memory address that is generated by the same step and needs software prefetching. All the work performed for a query can be modeled into a list of steps as defined here. Each thread gets *M* queries to execute at a time. On a single thread, if the queries are executed in sequential manner, each step of a query has to pay the memory latency cost in the memory access phase. Instead, we process a batch of *m* queries simultaneously.

**Figure 14:**
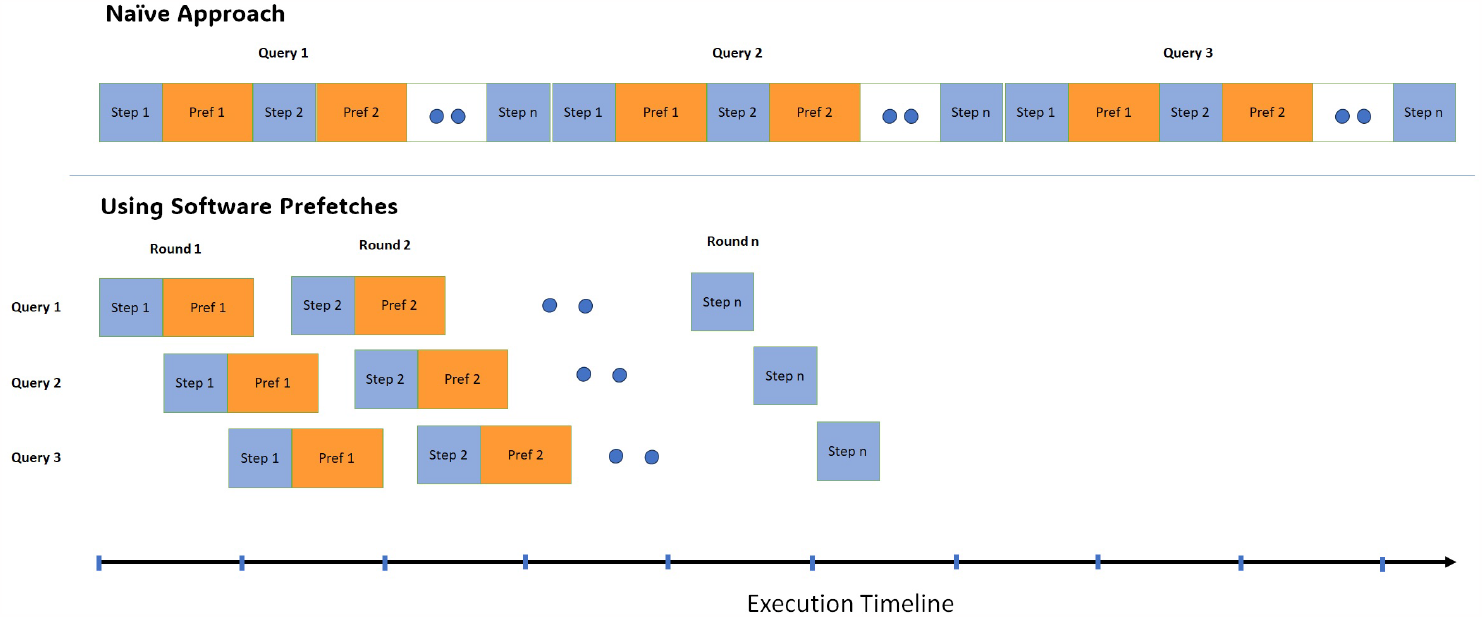
Illustration of our design for software prefetching. The top figure illustrates the naive approach which executes all the query processing steps before starting the next query. The bottom figure depicts the software prefetching approach with a batch of 3 queries executing the query processing steps in a round-robin fashion.

Let *q*_*i*_ denote the *i*-th query in a batch. We process the queries in a round robin fashion – processing one step at a time and prefetching for the next step. We first execute the setup step of *q*_0_ and start software prefetching the memory addresses for the next step. While the data is getting prefetched, we process the setup step of *q*_1_ and start its prefetching. We do this for all the queries in the batch and then return to *q*_0_ for the next step. By this time, data for next step of *q*_0_ is already in cache and we can perform the compute. We keep repeating this for all the steps for the queries. The queries may have different number of steps as in the case of binary search. When a query is done, we replace it with a new query to keep the batch size fixed. The batch size, *m*, should be big enough such that by the time we come back to a query, its data is already prefetched to cache. On the other hand, it should be small enough so that the data corresponding to all the queries in a batch can fit in cache. We theoretically calculate the range of batch sizes and then empirically find the best batch size near the theoretical range.

#### 5.12.3 Vectorization

The last mile search (Fig. 9b) is performed using a binary search. However, once we have less than 8 elements left to search, we compare with all of them simultaneouly using SIMD instructions.

#### 5.12.4 Multi-threading

We use a simple multi-threading design where we divide the set of queries into blocks of *M* queries and use dynamic scheduling to distribute the blocks of queries across threads.

#### 5.12.5 Benefit of Hardware-aware Optimizations

For these experiments, we use human genome as the reference sequence.

For query datasets, we use S1 and H1 (described in Supplementary Table 4 and 3) for *exact search* and *SMEM search*, respectively.

Supplementary Table 8 and 9 show the benefits of hardware-aware optimizations for *exact search* and *SMEM search*, respectively, on a single thread. We report performance of TAL and three versions of LISA: (1) without any hardware-aware optimizations (Un-opt), (2) with application of software prefetching to Un-opt (+SW pref.) and (3) with application of vectorization in addition to SW prefetching (+vectorization).

Compared to TAL, the number of instructions executed by our un-optimized implementations are 3.4× and 13× less, respectively, for *exact* and *SMEM search*. This shows the benefit of our algorithmic improvements to reduce the work required for sequence search. However, the un-optimized implementations suffer from significantly low L1 cache hit rate, resulting in poor throughput. Application of software prefetching to un-optimized LISA implementations dramatically improves the L1 hit rate, thus improving throughput, despite increasing the number of instructions. More specifically, software prefetching improves throughput by 3.36 and 3.86 for *exact search* and *SMEM search*, respectively. Through vectorization of the last few iterations of the binary search, we reduce the instructions to get a further 15% improvement in throughput for *exact search*. For *SMEM search*, the benefit of vectorization is only 2% because the last few iterations of binary search form only a small fraction of the total time. This shows that improvements in the algorithm and implementations that are well tuned to the hardware are both significant in LISA achieving high performance gains.

Supplementary Table 10 reports the memory bandwidth achieved by TAL and un-optimized and fully optimized versions of LISA on a full CPU socket. It is clear that our optimizations improve the bandwidth utilization of LISA. While LISA consumes significantly high memory bandwidth, given that it needs to read much less data from memory, it consumes significantly less time than TAL.

Comparing the performance of TAL with fully optimized LISA version, it is clear that the main benefit of LISA comes from reducing the work required to solve the problem, as evident in the reduced instruction counts and memory access requirements. In addition, the hardware-aware implementation ensures that the reduced amount of work translates to better performance.

## 6 RELATED WORK

### 6.1 Acceleration of FM-index based Search

A number of approaches have been proposed to accelerate FM-index based methods for modern multicore CPUs [8, 12, 13, 15, 22, 26, 29, 31, 32, 57], manycore CPUs [58, 59], GPUs [14, 27, 28, 60, 61], and FPGAs [30, 62]. A majority of them have targeted the simpler, exact search algorithm. We mention a few notable approaches here.

Compression of FM-index, first introduced in [25], is a widely used technique that reduces the memory requirement of FM-index. Compression of FM-index by, say a factor *h*, is achieved by storing the array *O* only for every *h*-th position and recomputing for the rest of the positions every time by using the closest position that is a multiple of *h* and the BWT. Compression also enables look-ups for more positions to hit in the same cache line, thereby, reducing the memory bandwidth requirement.

Trans-Omics Acceleration Library (TAL) [7, 32, 33] also uses a compressed FM-index and provides efficient hardware-aware implementations for CPUs. TAL demonstrates significant performance gains on *exact search, inexact search*, and *SMEM search* with application of software prefetching and vectorization on the FM-index based backward search algorithm, inexact search algorithm [15], and the *SMEM search* algorithm [45]. For *exact search* and *inexact search*, the software prefetching design used by TAL is similar to that presented in Sec. 5.12.

Chacon *et al*. [26, 27] developed a clever *n*-step FM-index strategy for *exact search* that allows processing *n* bases at a time. This is achieved by using Σ^*n*^ as alphabet instead of Σ. It reduces the number of memory accesses by a factor of *n* while increasing the number of executed instructions. Moreover, the memory requirement of this strategy is exponential in *n*, thus, only allowing *n*≤ 4 on most systems. They showed significant performance gains on CPU and GPGPU. However, their application of software prefetching on CPU for exact search yielded minimal benefit due to an inefficient design, resulting in their performance being memory latency bound.

For SMEM, [50, 54] use Enhanced Suffix Array to perform the shrink operation like us and uses FM-index for the extension. They lack the learned index based extension and efficient architecture-aware implementations of extend and shrink operations.

### 6.2 Learning-based Search

#### 6.2.1 Sapling

Sapling is a recently published learned index based solution for *exact search* that showed promising results [22]. Sapling is single threaded and does not provide architecture-aware optimizations. Given a query *Q*, Sapling outputs any one position in the BW-Matrix where the query matches instead of outputting the entire SA interval. Sapling learns a machine learning model for queries of fixed length *k*. Sapling paper proposes an artificial neural network (ANN) based model and a piece-wise linear model (PWL) and shows that PWL outperforms ANN. For the PWL model, Sapling divides the space of 4^*k*^ possible queries into equal sized intervals and divides the BW-Matrix into those intervals based on matching prefix. Within the interval, it uses a PWL model to predict the position of the query. It stores the maximum error for each interval to provide the bounds of last mile search (using binary search). The learned index used by Sapling is similar to the RadixSpline [41] with linear splines replaced with fixed length intervals. As mentioned in the Sapling paper, the fixed length intervals lead to very large bounds for binary search in some intervals. To alleviate that, Sapling also stores the 95th percentiles of errors to reduce the iterations of binary search. However, this still results in very large bounds for binary search. The best bound reported for Sapling is 653, which results in 10 iterations of binary search. RadixSpline resolves this by using linear splines to have a bound on the size of each interval. LISA uses RMI learned index structure that has empirically been shown to perform better than RadixSpline [41] and Piecewise Geometric Model (PGM) [36].

For queries with length less than *k*, they need to be appended by *A*’s to make their length *k*. For queries with length greater than *k*, if *v* is the integer representation of *Q* [0, *k* −1], it uses a floating point number between *v* and *v* +1 based on *Q* [*k*, |*Q*| −1] to evaluate the PWL function on. However, in the last mile search the entire length of the query needs to be compared in each step making it inefficient for long queries. On the other hand, LISA divides a query into chunks of size *K* and uses IPBWT to be able to efficiently handle queries of all lengths.

#### 6.22 bwa-meme

Both LISA and bwa-meme [23] apply learned approach to accelerate the sequence search (seeding) kernels of BWAMEM2 [7]. However, they follow completely different approaches. Bwa-meme maintains an array in which each entry represents a suffix of the reference sequence. Each entry consists of the first 32 bases of the suffix - represented using 64 bits with each base encoded using 2 bits - and the position of the suffix in the reference sequence. The array is sorted in lexicographic order of the suffixes. Bwa-meme trains a learned model on 32-base queries to search in this array. A query search is performed by using the learned model on the first 32 bases of the query to get an interval in the reference sequence. Subsequently, last mile search is performed through binary search in the interval. In each step of the binary search, the query is matched to the first 32 bases from the array. Longer queries beyond 32 bases are matched in the reference sequence.

Compared to LISA, the approach used by bwa-meme is unfavorable for longer queries. LISA uses IPBWT that supports the search of arbitrary long queries by searching for K bases at a time, thus using the learned model to narrow the search space for every K bases. On the other hand, bwa-meme performs the search of the entire query in one step applying the learned model only on the first 32 bases. For queries with highly frequent first 32 bases, the interval for last mile search of bwa-meme is long. Also, queries longer than 32 need to be matched in the reference sequence, which is slow. This is evident in the results shown in Section 2. LISA achieves around 1.07× -1.4× speedup over bwa-meme for the shorter reads of length 101 bases while it achieves around 1.4× -1.7× speedup over bwa-meme for longer reads with 250 bases.

## ACKNOWLEDGEMENTS

This research is supported by Google, Intel, and Microsoft as part of the MIT Data Systems and AI Lab (DSAIL) at MIT.

## Optimization Notice

Software and workloads used in performance tests may have been optimized for performance only on Intel microprocessors. Performance tests, such as SYSmark and MobileMark, are measured using specific computer systems, components, software, operations and functions. Any change to any of those factors may cause the results to vary. You should consult other information and performance tests to assist you in fully evaluating your contemplated purchases, including the performance of that product when combined with other products. For more information go to http://www.intel.com/performance. Intel, Xeon, and Intel Xeon Phi are trademarks of Intel Corporation in the U.S. and/or other countries.

## A SUPPLEMENTARY

### A.1 Experimental Setup

**Supplementary Table 1:**
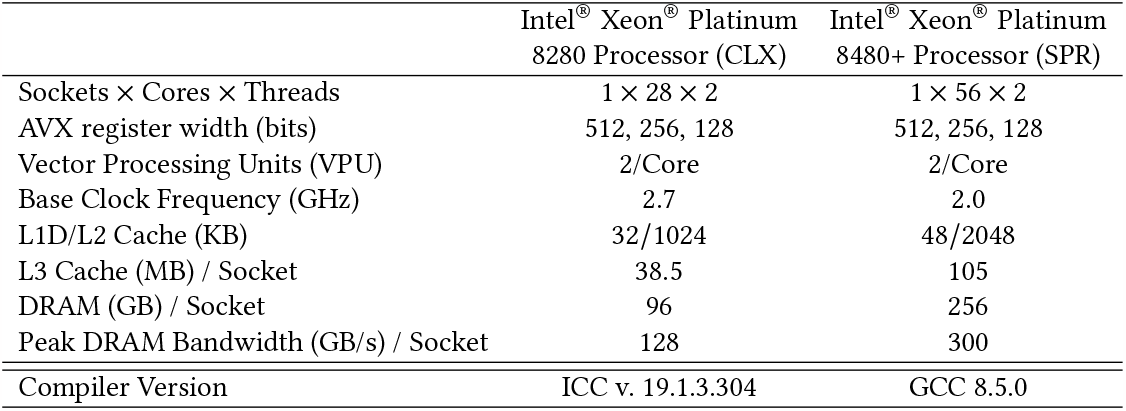
System Configuration.

|  | Intel® Xeon® Platinum<br>8280 Processor (CLX) | Intel® Xeon® Platinum<br>8480+ Processor (SPR) |
| --- | --- | --- |
| Sockets × Cores × Threads | $1 \times 28 \times 2$ | $1 \times 56 \times 2$ |
| AVX register width (bits) | 512, 256, 128 | 512, 256, 128 |
| Vector Processing Units (VPU) | 2/Core | 2/Core |
| Base Clock Frequency (GHz) | 2.7 | 2.0 |
| L1D/L2 Cache (KB) | 32/1024 | 48/2048 |
| L3 Cache (MB) / Socket | 38.5 | 105 |
| DRAM (GB) / Socket | 96 | 256 |
| Peak DRAM Bandwidth (GB/s) / Socket | 128 | 300 |
| Compiler Version | ICC v. 19.1.3.304 | GCC 8.5.0 |

**Supplementary Table 2:** Reference Sequences.

| Reference Sequence | Length<br>(Million bases) | Version |
| --- | --- | --- |
| Human | 3101 | hs37d5 |
| Asian Rice | 387 | IR64 (IRRI) |
| Zebra Fish | 1679 | GRCz11 |

**Supplementary Table 3:** Read datasets used for *SMEM search* experiments. In each case, we sampled the first 5 million reads from the source dataset.

| Read<br>Datasets | Organism | Length | No. of<br>Reads | Source<br>(NCBI-SRA) |
| --- | --- | --- | --- | --- |
| H1 | Human | 151 | 5 million | ERR2990063 |
| H2 | Human | 151 | 5 million | ERR3239330 |
| H3 | Human | 151 | 5 million | SRR7733443 |
| H4 | Human | 101 | 5 million | SRR622461 |
| H5 | Human | 101 | 5 million | SRR622457 |
| A1 | Asian Rice | 151 | 5 million | SRR10724346 |
| A2 | Asian Rice | 151 | 5 million | SRR10838753 |
| A3 | Asian Rice | 151 | 5 million | SRR8241153 |
| Z1 | Zebra Fish | 151 | 5 million | ERR2624531 |
| Z2 | Zebra Fish | 151 | 5 million | ERR3333446 |
| Z3 | Zebra Fish | 151 | 5 million | SRR10958316 |

**Supplementary Table 4:** Seed datasets used for *exact search* experiments. We sample fixed length seeds from the real original read datasets.

| Seed Datasets | Length | No. of Seeds | Original Read Dataset (NCBI-SRA). |
| --- | --- | --- | --- |
| S1 | 22 | 50 million | ERR2990063 |
| S2 | 22 | 50 million | ERR3239330 |
| S3 | 22 | 50 million | SRR7733443 |
| S4 | 22 | 50 million | SRR622461 |
| S5 | 22 | 50 million | SRR622457 |
| S6 | 22 | 50 million | SRR10724346 |
| S7 | 22 | 50 million | SRR10838753 |
| S8 | 22 | 50 million | SRR8241153 |
| S9 | 22 | 50 million | ERR2624531 |
| S10 | 22 | 50 million | ERR3333446 |
| S11 | 22 | 50 million | SRR10958316 |

**Supplementary Table 5:**
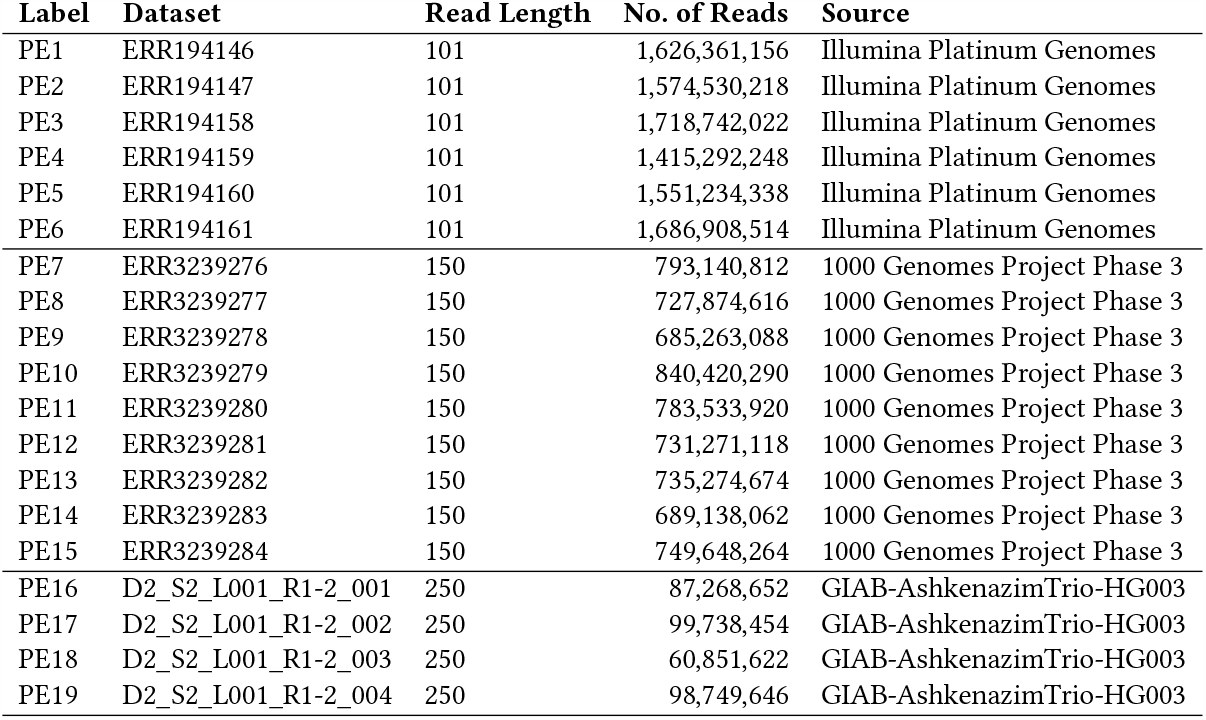
The paired-end (PE) short read datasets (PE1 to PE19) used for bwa-mem2, bwa-meme and LISA comparison. Each dataset consists of two files: R1 and R2 containing the exact same number of reads.

| Label | Dataset | Read Length | No. of Reads | Source |
| --- | --- | --- | --- | --- |
| PE1 | ERR194146 | 101 | 1,626,361,156 | Illumina Platinum Genomes |
| PE2 | ERR194147 | 101 | 1,574,530,218 | Illumina Platinum Genomes |
| PE3 | ERR194158 | 101 | 1,718,742,022 | Illumina Platinum Genomes |
| PE4 | ERR194159 | 101 | 1,415,292,248 | Illumina Platinum Genomes |
| PE5 | ERR194160 | 101 | 1,551,234,338 | Illumina Platinum Genomes |
| PE6 | ERR194161 | 101 | 1,686,908,514 | Illumina Platinum Genomes |
| PE7 | ERR3239276 | 150 | 793,140,812 | 1000 Genomes Project Phase 3 |
| PE8 | ERR3239277 | 150 | 727,874,616 | 1000 Genomes Project Phase 3 |
| PE9 | ERR3239278 | 150 | 685,263,088 | 1000 Genomes Project Phase 3 |
| PE10 | ERR3239279 | 150 | 840,420,290 | 1000 Genomes Project Phase 3 |
| PE11 | ERR3239280 | 150 | 783,533,920 | 1000 Genomes Project Phase 3 |
| PE12 | ERR3239281 | 150 | 731,271,118 | 1000 Genomes Project Phase 3 |
| PE13 | ERR3239282 | 150 | 735,274,674 | 1000 Genomes Project Phase 3 |
| PE14 | ERR3239283 | 150 | 689,138,062 | 1000 Genomes Project Phase 3 |
| PE15 | ERR3239284 | 150 | 749,648,264 | 1000 Genomes Project Phase 3 |
| PE16 | D2_S2_L001_R1-2_001 | 250 | 87,268,652 | GIAB-AshkenazimTrio-HG003 |
| PE17 | D2_S2_L001_R1-2_002 | 250 | 99,738,454 | GIAB-AshkenazimTrio-HG003 |
| PE18 | D2_S2_L001_R1-2_003 | 250 | 60,851,622 | GIAB-AshkenazimTrio-HG003 |
| PE19 | D2_S2_L001_R1-2_004 | 250 | 98,749,646 | GIAB-AshkenazimTrio-HG003 |

### A.2 Algorithms

#### Supplementary Algorithm 1

Exact Search Algorithm using FM-index

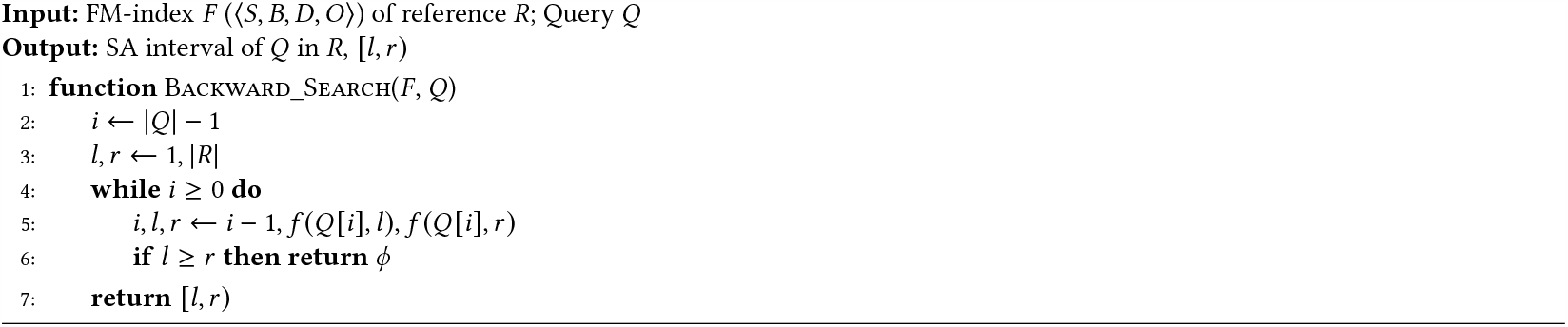

#### Supplementary Algorithm 2

Exact-LISA: Exact Search Algorithm using IPBWT

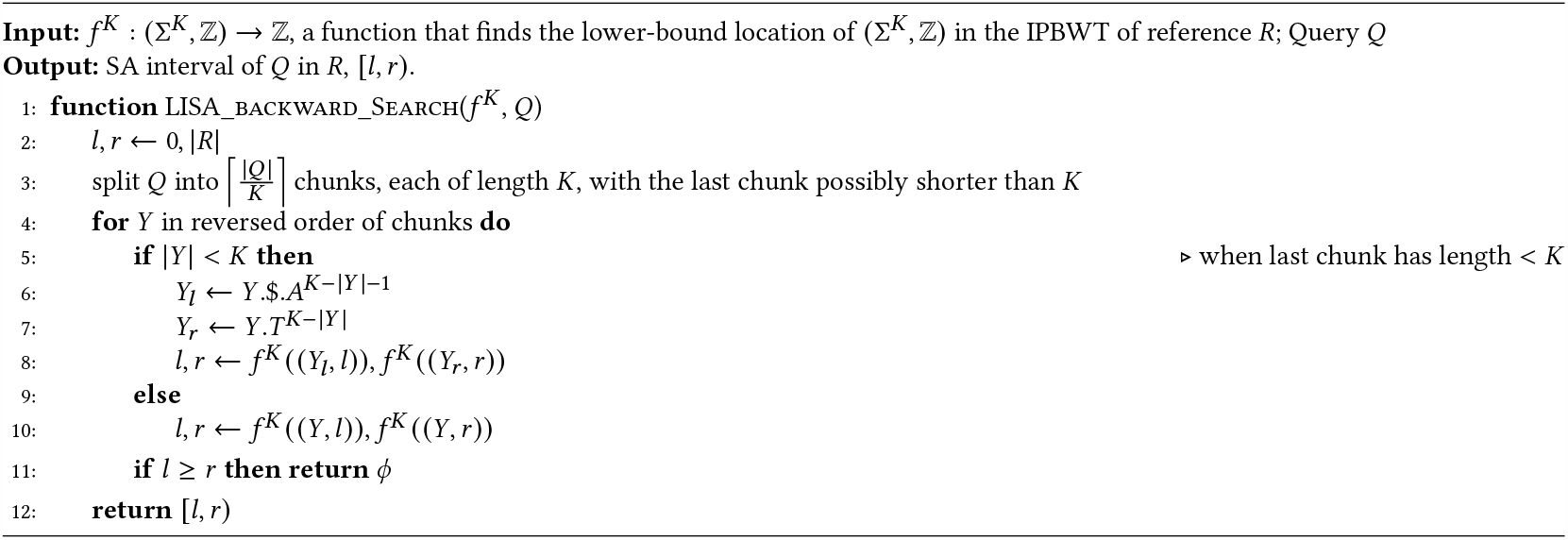

#### Supplementary Algorithm 3

SMEM Algorithm using Extend and Shrink

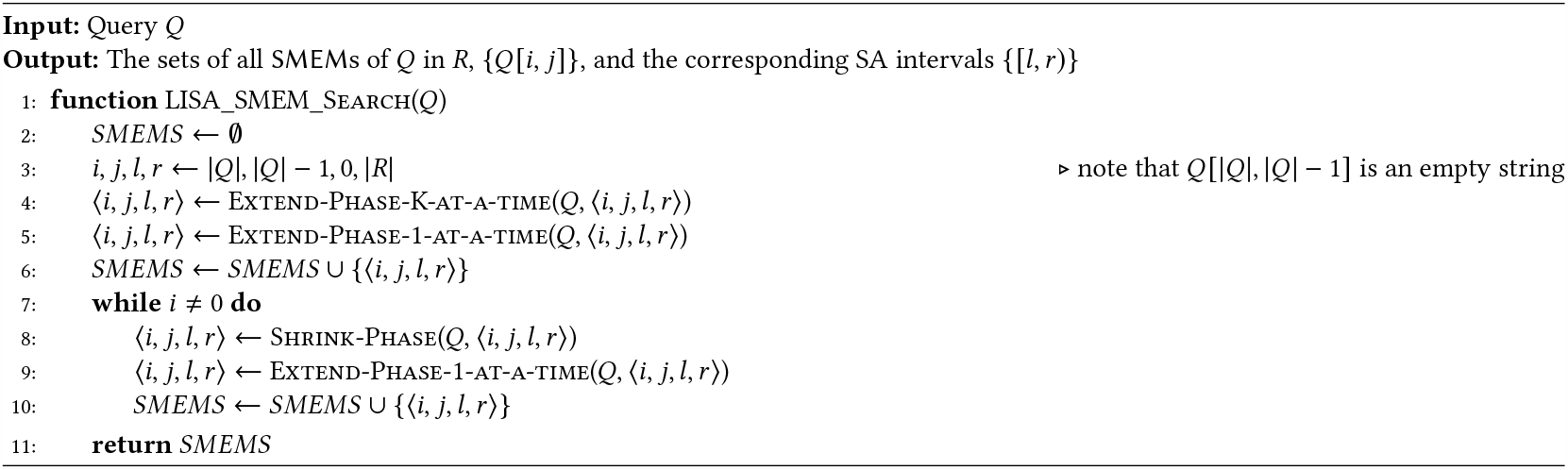

#### Supplementary Algorithm 4

Algorithm for extending one base at a time

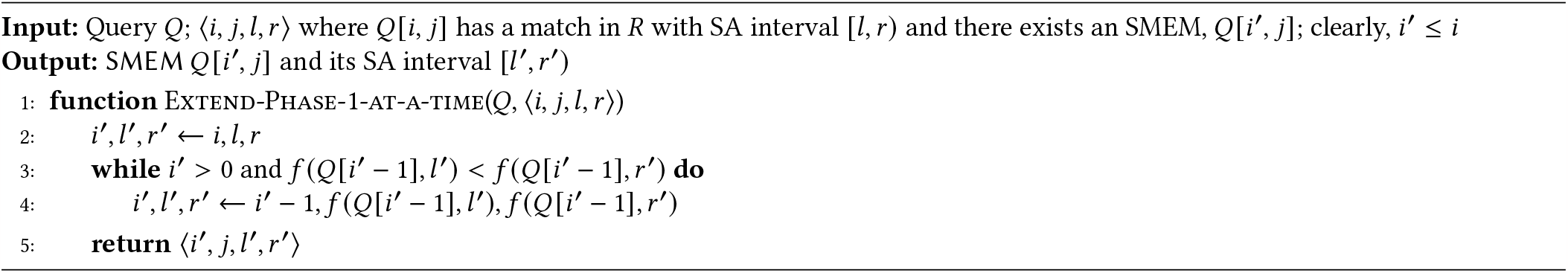

#### Supplementary Algorithm 5

Algorithm for extending *K* bases at a time

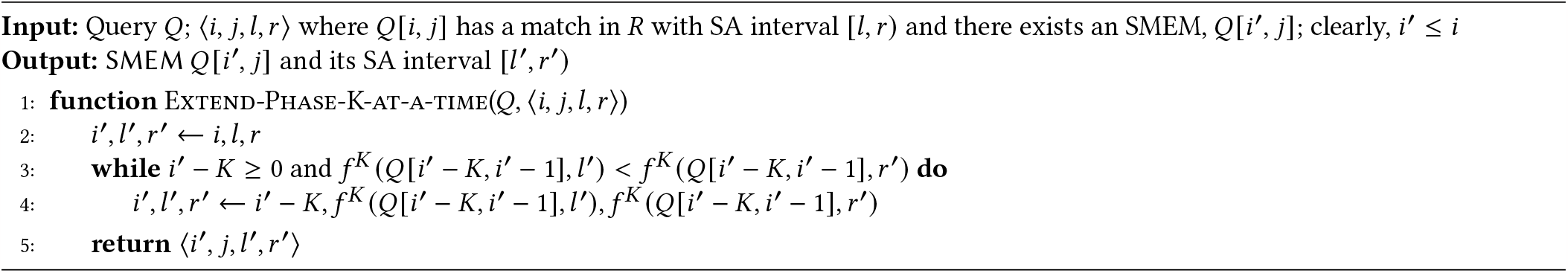

#### Supplementary Algorithm 6

Algorithm for one Shrink Operation

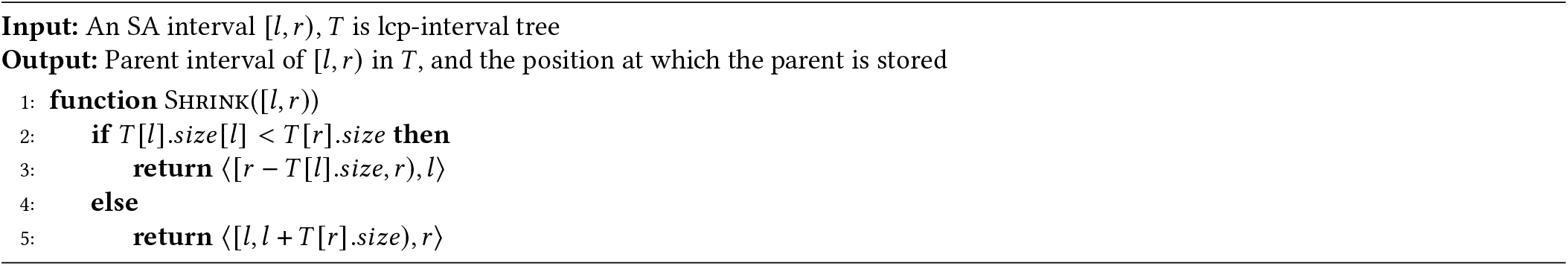

#### Supplementary Algorithm 7

Proposed Algorithm for the Shrink Phase

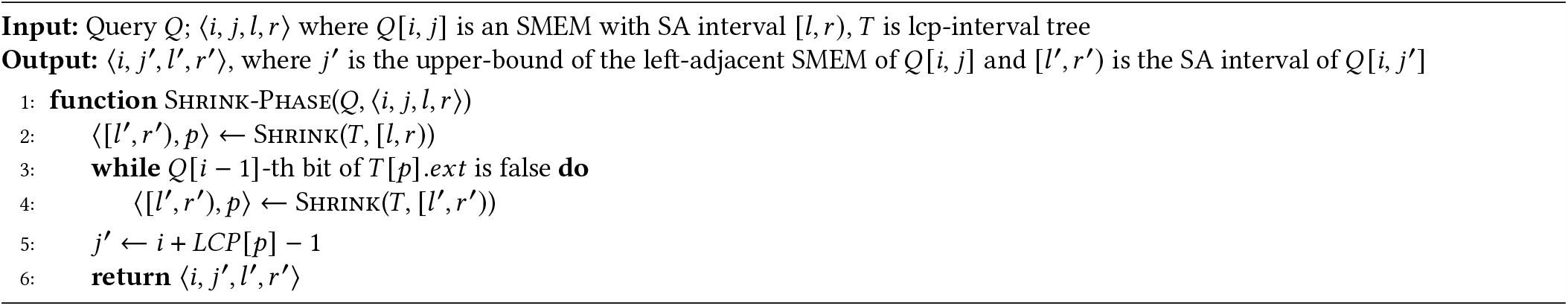

#### Supplementary Algorithm 8

Proposed algorithm for BWA-MEM2: SMEM using Extend and Shrink

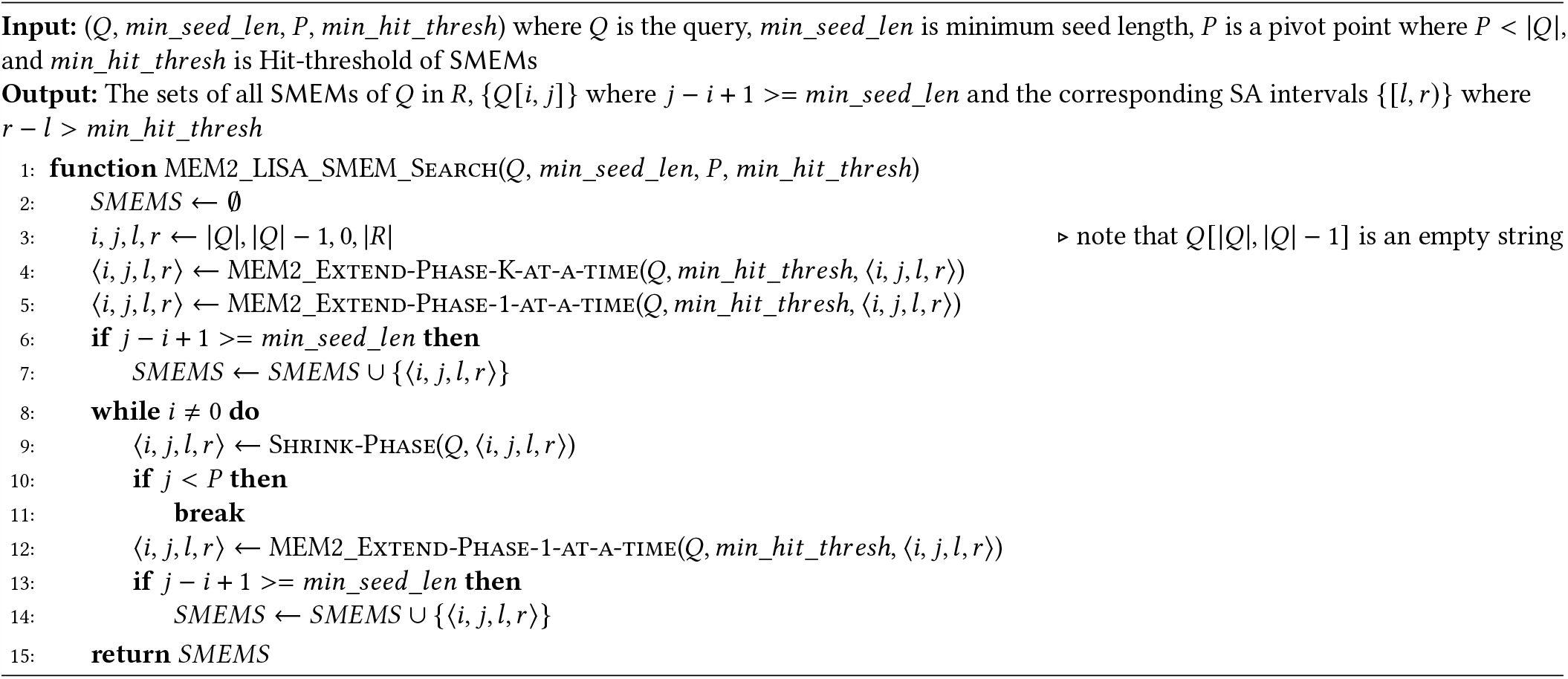

#### Supplementary Algorithm 9

Proposed algorithm for BWA-MEM2: extend one base at a time

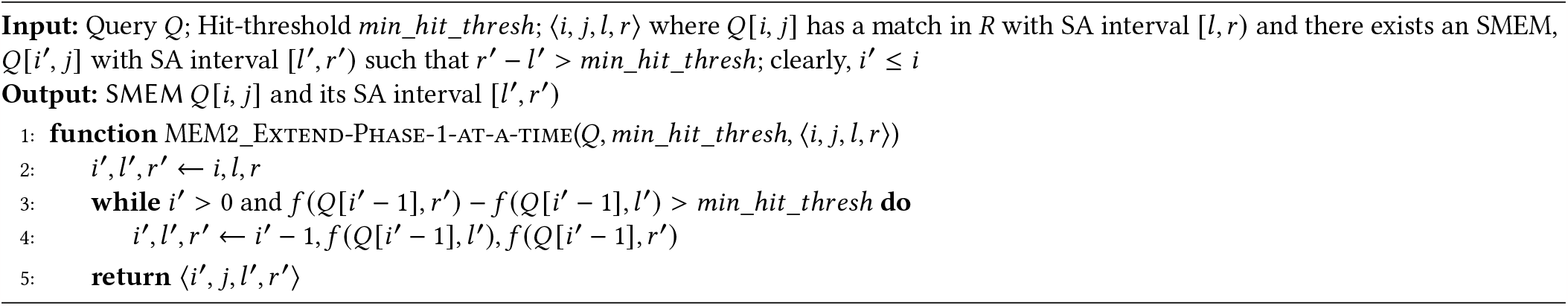

#### Supplementary Algorithm 10

Proposed algorithm for BWA-MEM2: extend *K* bases at a time

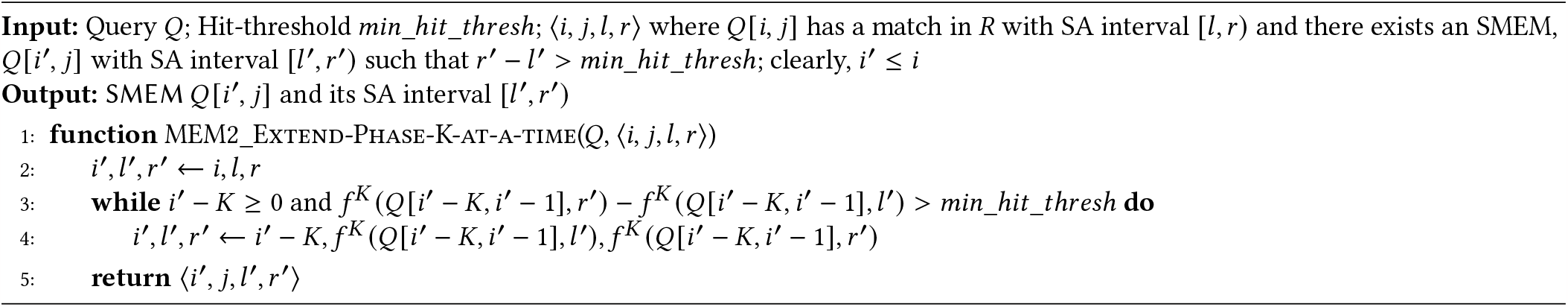

#### Supplementary Algorithm 11

Proposed algorithm for BWA-MEM2: LISA-based Seed Strategy

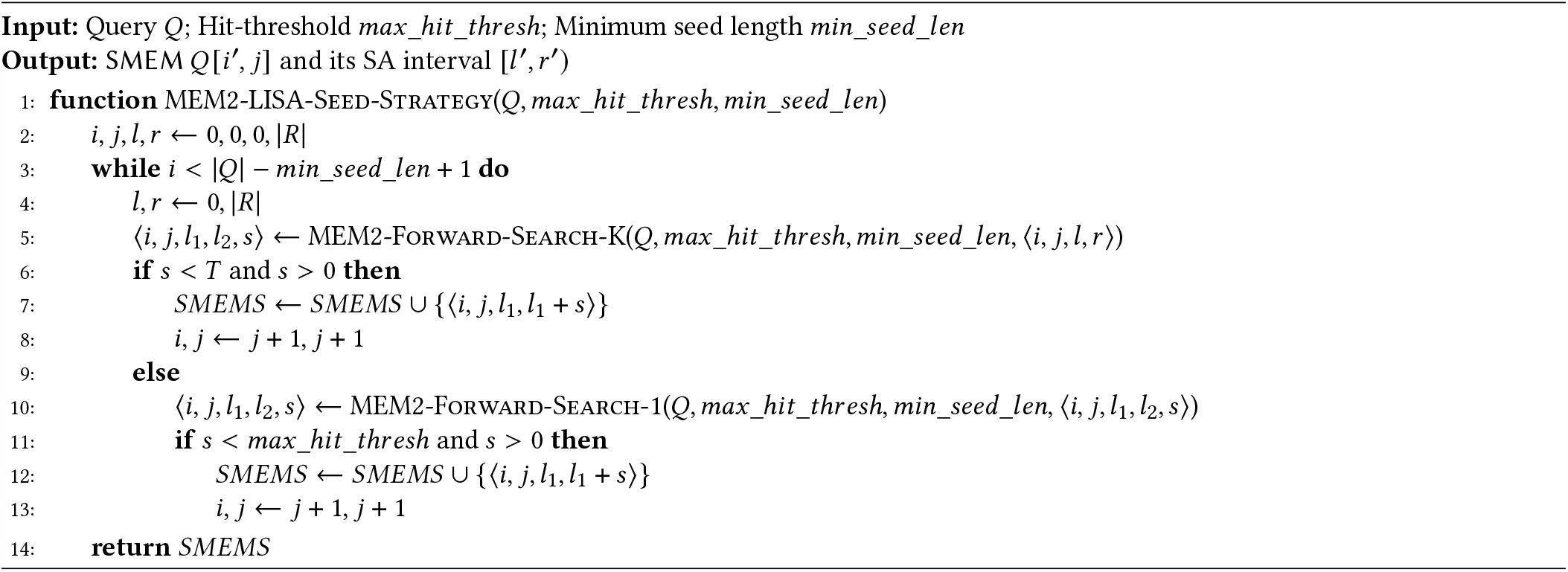

#### Supplementary Algorithm 12

Proposed algorithm for BWA-MEM2: forward exact-search of *K* bases

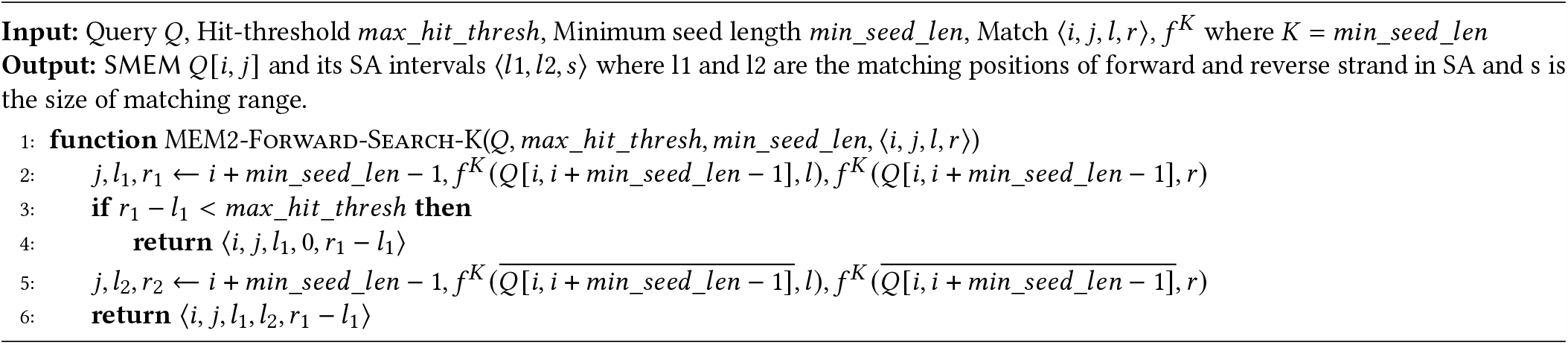

#### Supplementary Algorithm 13

Proposed algorithm for BWA-MEM2: forward extension by one base at a time

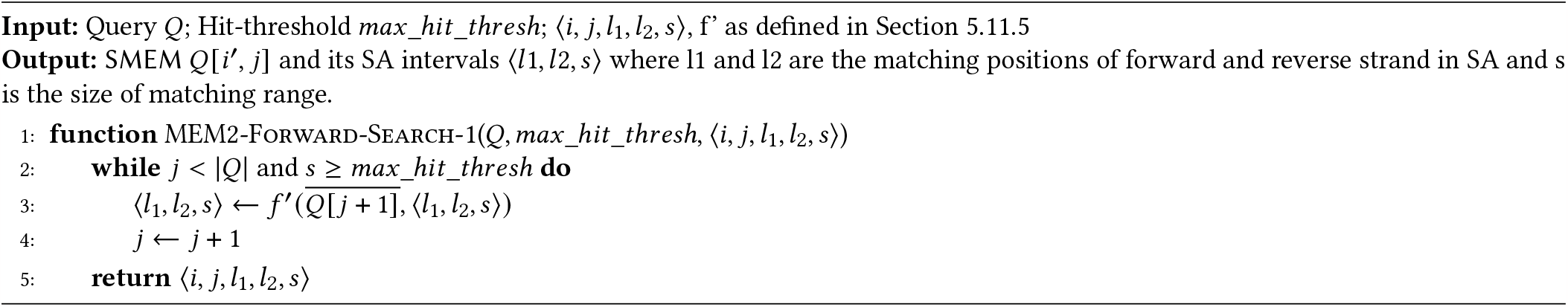

#### Supplementary Algorithm 14

Proposed algorithm for BWA-MEM2: LISA-based Seeding

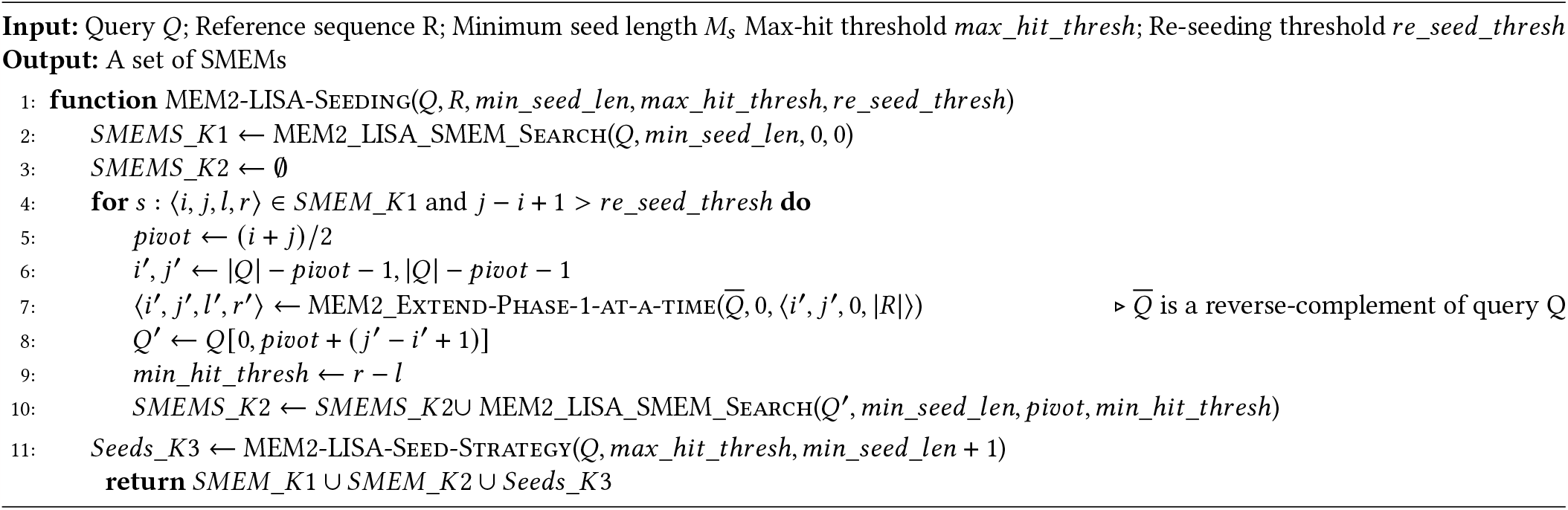

### A.3 Additional Results

**Supplementary Figure 1:**
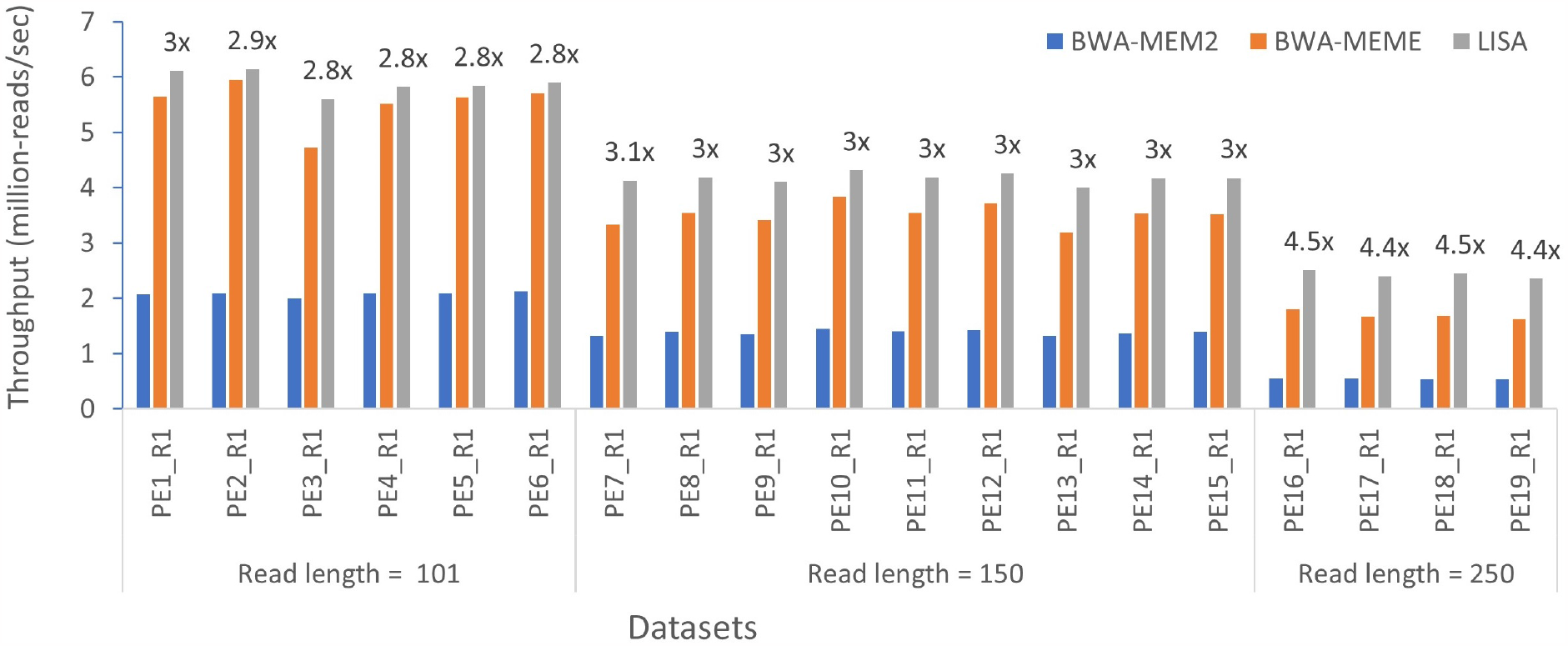
Performance evaluation for *seeding phase* of BWA-MEM2 on a single CLX CPU (28 cores with 56 threads) for single-ended read datasets. Throughput gain of LISA over BWA-MEM2 is shown on the top of LISA bars.

**Supplementary Table 6:** Effect of number of RMI leaf nodes. For this experiment, we use synthetic seeds of length 22 that are sampled from the corresponding reference genome sequence. We report the throughput in Million Seeds Per Second (MSPS), average number of binary search steps per last-mile search and memory footprint of RMI with respect to the increase in the number of RMI leaf nodes.

| Reference genome | # RMI leaves | Throughput (MSPS) | # Binary search steps | Memory footprint (GB) |
| --- | --- | --- | --- | --- |
| **Human** | $2^{25}$ | 10.4 | 9.17 | 0.8 |
| | $2^{26}$ | 11.2 | 7.88 | 1.6 |
| | $2^{27}$ | 11.7 | 7.00 | 3.2 |
| | $2^{28}$ | 12.1 | 6.00 | 6.1 |
| | $2^{29}$ | 12.4 | 5.46 | 13 |
| **Asian Rice** | $2^{21}$ | 10.9 | 8.67 | 0.05 |
| | $2^{22}$ | 12.4 | 7.29 | 0.10 |
| | $2^{23}$ | 13.2 | 6.18 | 0.20 |
| | $2^{24}$ | 13.5 | 5.14 | 0.40 |
| | $2^{25}$ | 14.3 | 4.41 | 0.80 |
| **Zebra Fish** | $2^{24}$ | 10.3 | 9.57 | 0.40 |
| | $2^{25}$ | 10.7 | 8.55 | 0.80 |
| | $2^{26}$ | 11.3 | 7.47 | 1.6 |
| | $2^{27}$ | 11.7 | 6.70 | 3.1 |
| | $2^{28}$ | 12.3 | 5.91 | 6.1 |

**Supplementary Table 7:**
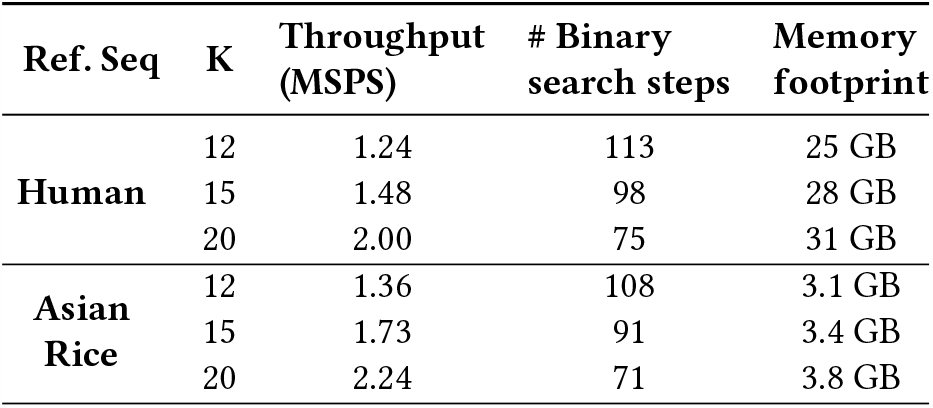
Effect of K. For this experiment, we compare three values of *K* (12, 15 and 20) by using synthetic seeds of length 120 (which is divisible by all three values) that are sampled from the corresponding reference genome sequence. We report the throughput in Million Seeds Per Second (MSPS), average number of binary search steps per seed and memory footprint of IPBWT with respect to the increase in the value of *K* .

**Supplementary Figure 2:**
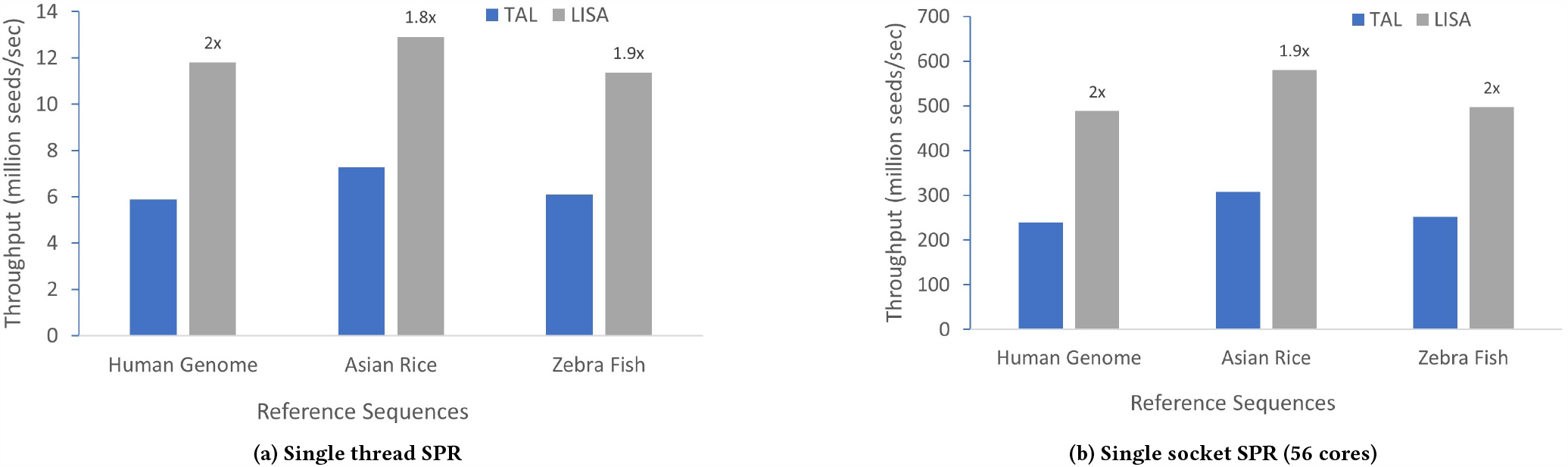
Performance evaluation for *exact search* problem on synthetic queries. For each experiment, we extracted 50 Million synthetic seeds of length 22 from the corresponding genomes. The speedup of LISA over TAL is shown at the top of each LISA bar. *K* = 22 and # of leaf nodes is 2^27^, 2^23^ and 2^26^ for Human, Asian Rice and Zebra Fish, respectively.

**Supplementary Figure 3:**
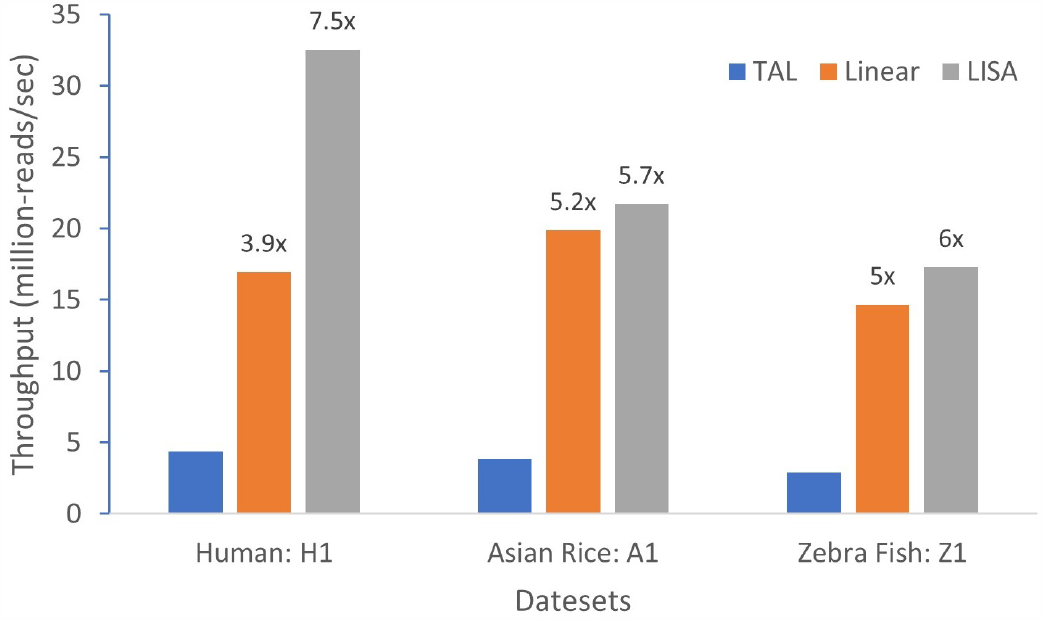
Performance comparison of TAL, our ESA based O(|Q|) SMEM algorithm (Linear) and SMEM-LISA algorithm (LISA) for *SMEM search* problem using a single socket (56 cores) of SPR CPU. We use Human, Asian Rice and Zebra Fish genomes as reference sequences and H1, A1 and Z1 as read datasets. The speedup of LISA/Linear over TAL is shown at the top of each LISA/Linear bar. *K* = 22 and # of leaf nodes is 2^28^, 2^24^ and 2^27^ for Human, Asian Rice and Zebra Fish, respectively.

**Supplementary Figure 4:**
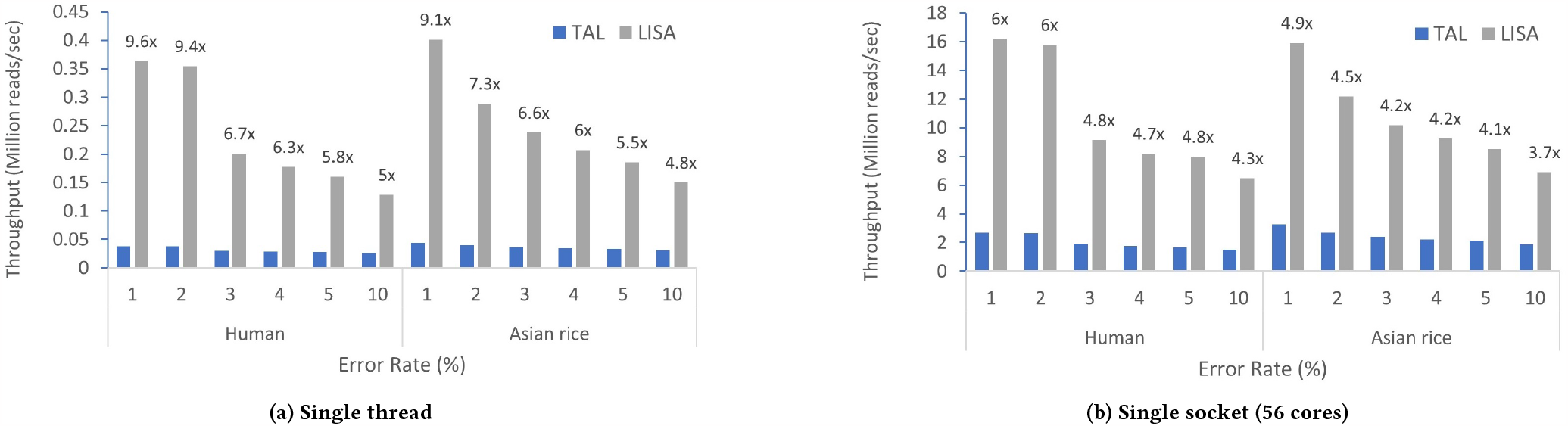
Effect of varying error rate in the read datasets on throughput for *SMEM search* problem. For each experiment, 5 Million synthetic reads of length 151 were generated using wgsim [56]. The speedup of LISA over TAL is shown at the top of each LISA bar. *K* = 22 and # of leaf nodes is 2^28^ and 2^24^ for Human and Asian Rice, respectively.

**Supplementary Table 8:** Effect of hardware-aware optimizations for *exact search* on a single thread. Reference sequence: Human Genome. Query dataset: S1. We report throughput in Million Seeds Per Second (MSPS), instruction count and percentage of loads hitting in L1 cache (L1 hit rate) for TAL and three versions of LISA.

|  | TAL | LISA |  |  |
| --- | --- | --- | --- | --- |
|  |  | Un-opt | +SW pref. | +vectorization |
| **Throughput (MSPS)** | 4.49 | 2.29 | 7.69 | 8.82 |
| **Inst. count** | 9.2E+10 | 2.7E+10 | 4.1E+10 | 3E+10 |
| **L1 hit rate** | 99.6% | 85% | 99.6% | 99.6% |

**Supplementary Table 9:** Effect of hardware-aware optimizations for *SMEM search* on a single thread. Reference sequence: Human Genome. Query dataset: S1. We report throughput in Million Reads Per Second (MRPS), instruction count and percentage of loads hitting in L1 cache (L1 hit rate) for TAL and three versions of LISA.

|  | TAL | LISA |  |  |
| --- | --- | --- | --- | --- |
|  |  | Un-opt | +SW pref. | +vectorization |
| **Throughput (MRPS)** | 0.04 | 0.14 | 0.54 | 0.55 |
| **Inst. count** | 3.9E+11 | 3E+10 | 4.4E+10 | 3.8E+10 |
| **L1 hit rate** | 91% | 80.7% | 97.7% | 98.2% |

**Supplementary Table 10:** Memory bandwidth utilization (GB/s) on a single CLX CPU socket (28 cores) for *exact search* and *SMEM search* for TAL and two versions of LISA.

|  | TAL | LISA Un-opt | LISA Optimized |
| --- | --- | --- | --- |
| **Exact search** | 79.6 | 15 | 78.2 |
| **SMEM search** | 21.5 | 24 | 65.8 |

### A.4 Sapling vs TAL vs LISA

**Supplementary Figure 5:**
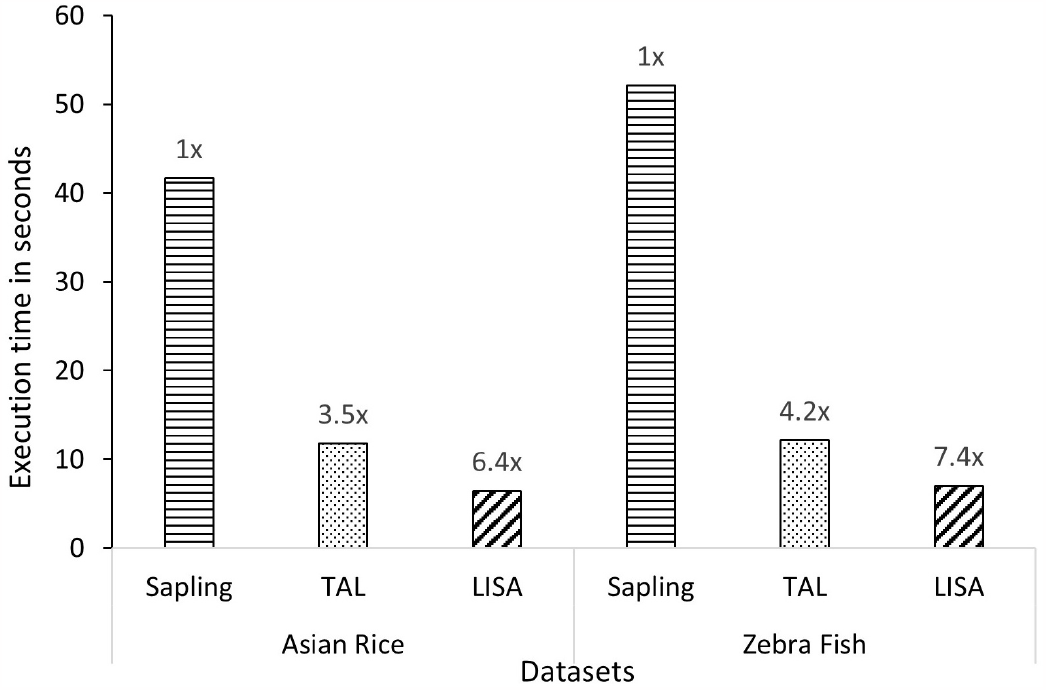
Performance of the *exact search* for Sapling, TAL, and LISA using a single thread.

Here, we compare the execution time of LISA and TAL with the recently published Sapling for *exact search* [22]. Sapling demonstrated a speedup of over 2× over Bowtie, Mummer, and an optimized implementation of binary search. Therefore, we omit any comparison with Bowtie, Mummer, and binary search. Moreover, Sapling is single threaded. Therefore, we compare the performance of the three implementations using only a single thread. We have used the same evaluation method as used in the Sapling paper. We use the scripts provided with the Sapling source code to generate 50 million seeds of length 21 for the three reference sequences [63]. The script ensures that there is at least one match of the generated seeds.

Supplementary Fig. 5 shows the comparison. Note that the time reported for Sapling here is more than 2× less than that reported in [22] potentially due to a difference in the CPU used. Moreover, Sapling does not return the full interval in the suffix array in which the query matches, but rather returns one position in the interval – one needs to perform a search on both sides of the position to get the interval. So, the time required for Sapling based method for realistic usage would be higher than reported here. We could not run Sapling for human genome as it ran out of memory even when run on a different machine with 256 GB DRAM. The time reported for TAL and LISA is the time spent in getting the full interval.

Supplementary Fig. 5 shows that TAL is 3.5× and 4.2× faster than Sapling and LISA is 6.4× and 7.4× faster than Sapling, respectively, for

Asian Rice and Zebra Fish. Therefore, for the rest of the paper, we only compare LISA with TAL.

### A.5 Time Complexity of the SMEM-LISA Algorithm

For this analysis, let us first assume that we do not use learned approach to extend *K* bases at a time, but instead use FM-index based 1 base at a time extension. Therefore, for extend phase, every decrement of *i* ^′^ (Supplementary Alg. 4, line 4) takes *O* 1 time. As for shrink phase, each shrink operation (Supplementary Alg. 7, line 4) takes *O* (1) time. If there were no dummy nodes, the structure of suffix tree guarantees that *j* ^′^ decreases by at least 1 per shrink operation. After dummy nodes are inserted, because each true node can have at most |{*A, C, G, T*, $}| = 5 children, parents and children are separated by at most 5 or *O* (1) dummy nodes. So, it takes at most *O* (1) extra calls for shrink to reach the true parent. Hence the time complexity of each decrement of *j* ^′^is *O* (1). There can be at most |*Q* | decrements to *i* ^′^ and *j* ^′^, so the total time complexity is *O* (|*Q* |).

When we use learned approach for the first phase of extension, the worst case complexity of an extension by *K* bases is *O* (*log* |*R* |) . This is because we need to perform binary search on the interval returned by the RMI, which could be the entire IPBWT in the worst case if we have a terrible model. However, as shown for exact search, extension by *K* using learned approach is faster than extension by *K* letters using FM-index, thus achieving lower than O(*K*) complexity in practice. Therefore, our LISA based SMEM algorithm is sub linear in |*Q* | in practice.

## Notes

### Competing Interest Statement

The authors have declared no competing interest.

### Summary of Updates

In this version, we make the following changes: 1. We make a case for wider usage of LISA. We do this by demonstrating LISA's applicability in additional use-cases. For example, we demonstrate applicability of LISA for accelerating the seeding kernels of BWA-MEM2. Section 5.11 discusses the design of LISA-based seeding kernels in BWA-MEM2. Figure 4 and Supplementary-Figure 1 present the performance comparison. Algorithms are added to the supplementary section. We also discuss how LISA has been applied to minimap2 and can also be applied to other algorithms. 2. A new set of results for exact search and SMEM search running on Intel Xeon Platinum 8480+ processor are added to the paper. 3. More figures are added to improve readability.

